# *Staphylococcus aureus* toxin LukSF dissociates from its membrane receptor target to enable renewed ligand sequestration

**DOI:** 10.1101/251645

**Authors:** Karita Haapasalo, Adam J. M. Wollman, Carla de Haas, Kok van Kessel, Jos van Strijp, Mark C. Leake

## Abstract

**background:** *Staphylococcus aureus* Panton Valentine Leukocidin (PVL) is a pore-forming toxin targeting the human C5a receptor (hC5aR), enabling this pathogen to battle the immune response by destroying phagocytes through targeted lysis. The mechanisms that contribute to rapid cell lysis are largely unexplored.

**Results:** Here we show that cell lysis may be enabled by a process of toxins targeting receptor clusters and receptor ‘recycling’ which allows multiple toxin pores to be formed close together. Using live cell single-molecule super-resolution imaging, Förster resonance energy transfer (FRET) and nanoscale total internal reflection fluorescence (TIRF) colocalization microscopy we visualized toxin pore formation in the presence of its natural docking ligand.

**Conclusions:** We demonstrate disassociation of hC5aR from toxin complexes and simultaneous binding of new ligands. This effect may free mobile receptors to amplify hyper inflammatory reactions in early stages of microbial infections and have implications for several other similar bi-component toxins and the design of new antibiotics.

## BACKGROUND

*S. aureus* causes diseases ranging from superficial skin and soft tissue infections (SSTI) to severe invasive diseases like osteomyelitis and necrotizing pneumonia [1]. During the 1960s, methicillin-resistant *Staphylococcus aureus* (MRSA) was identified as a nosocomial pathogen [2]. In the 1990s, infection of previously healthy community-dwelling individuals with MRSA was reported [3]. Since then, these community-associated (CA-) MRSA have rapidly emerged worldwide [4]. Variants have also recently been identified which have reduced susceptibility to the antibiotic vancomycin [5] as well as complete resistance, so-called VRSA [6], and these forms of *S. aureus* pose a significant threat to human health. S. *aureus* and resistant variants have also evolved adaptations to evade attack from cells of the human immune system. However, the molecular processes which underlie these strategies are underexplored in living cells. There are compelling scientific and societal motivations to understand the mechanisms involved in immunogenic evasion strategies of *S. aureus*.

In the early 1930s, Panton and Valentine described a powerful leukocidal toxin produced by multiple *S. aureus* isolates, now denoted Panton-Valentine Leukocidin (PVL), years later shown to be cytotoxic to neutrophils, monocytes and macrophages but not to lymphocytes [7, 8]. The majority of CA-MRSA isolates carry the genes encoding PVL, partially due to the successful spread of the PVL carrying clone USA300 in the USA [3, 4, 9, 10], rarely present in hospital-acquired antimicrobial resistant MRSA and methicillin susceptible *S. aureus* isolates. Based on epidemiological studies, PVL is associated with primary skin infections in humans, osteomyelitis and, in particular, severe necrotizing pneumonia [11, 12]. Necrotizing pneumonia is a severe complication caused by bacterial lung infection. It is characterized by massive recruitment of neutrophils in the site of infection, diffuse pulmonary inflammation, septic shock, and respiratory failure. Both host factors and microbial virulence factors are thought to play an important role in the inflammation, however, it is unknown how the interplay between these two factors affects the severity of the disease [13]. The specificity to cell surface receptors makes it difficult to study PVL’s role in *S. aureus* pathogenesis in a whole animal model. It is possible that lysis of neutrophils by PVL is responsible for a reduced host defense response allowing the pathogen to spread and cause eventual tissue damage. However, a previous study using a rabbit animal model on necrotizing pneumonia suggests that PVL itself directly or indirectly causes tissue injury and by this way induces local inflammation [14].

PVL is a pro-phage encoded bi-component, β-barrel pore-forming toxin (β-PFT) comprising protein subunits LukS and LukF. LukS binding to the surface of target cells induces secondary LukF binding; chemical and genetic analysis suggests that the resulting complex consists of a lytic pore-forming hetero-octamer [15, 16]. Stoichiometric analysis *in vitro* of this complex suggests it is an octamer of 4-plus-4 subunits [17]. In this complex only LukS is known to interact with the human C5a receptor (hC5aR, CD88), a G-protein coupled seven-transmembrane receptor (GPCR). LukS targets at least the extracellular N-terminus of hC5aR [18, 19], similar to the chemotaxis inhibitory protein of *S. aureus* (CHIPS), but may also interact with the transmembrane receptor region [20]. C5aR is the ligand for C5a, a powerful anaphylatoxin released during complement activation. Complement is a powerful first line defense mechanism against invading pathogens which can be initiated through three pathways: the classical, lectin, or alternative pathways. Activation of any of the three pathways on the target leads to a rapid opsonization with C3b [21]. Further activation of complement leads to initiation of the terminal pathway with release of C5a and formation of membrane attack complexes that are lytic for Gram-negative but not Gram-positive bacteria [22, 23]. Therefore, in defense against Gram-positive bacteria C3b-opsonization together with attraction and activation of neutrophils via C5a-C5aR interaction are essential [24, 25]. In severe cases, formation of C5a can potentially lead to hyper activation of the inflammatory response, an inability to regulate this potentially fatal reaction and eventually harm the human host tissues. Because of this strong pro-inflammatory activity, therapeutic interventions have recently focused on neutralizing antibodies against C5a and C5aR as potential candidates for the treatment of severe inflammatory conditions such as bacterial induced sepsis [26, 27].

LukS binding to hC5aR inhibits C5a receptor binding which efficiently blocks neutrophil activation [18]. LukS receptor binding alone is not sufficient for cell lysis but requires simultaneous interaction between the leukocidin subunits and hC5aR. However, multiple possible subunit and receptor combinations are theoretically possible and the spatiotemporal dynamics in functional complexes in live cells between LukS, LukF and hC5aR is not yet known. In addition to PVL *S. aureus* can produce a number of other β-PFTs with varying receptor and cell type specificities. From these LukED, LukAB (or LukGH) and γ-hemolysin (composed of two compound pairs, HlgA/HlgB or HlgC/HlgB) are classified as bi-component toxins like PVL while α-hemolysin is the prototypical β-PFT that assembles into a pore through the oligomerization of seven monomeric polypeptides [28].

Next to bacterial toxins, an entire group of other pore forming proteins have been identified which are involved in human innate immunity, indicating that pore-forming proteins are employed in survival strategies for several types of organisms [29]. Development of methods to study dynamic processes of pore formation by these toxins at a molecular level may improve our understanding of the evolution of bacterial virulence and human immunity. There are several studies that have attempted to explain the function of bacterial PFTs, including structural and subunit stoichiometry data from high resolution X-ray crystallography and single-molecule fluorescence microscopy [17, 30, 31]. However, these studies focused on pathogen instead of host factors and were thereby limited in excluding the specific interaction between host cell receptor and bacterial toxin component, the first step required for toxin oligomerization on the host cell membrane and the presence of the most potent factor mediating the inflammatory response via C5a recognition in the site of infection [18].

Here, we used standard and single-molecule fluorescence detection with super-resolution localization microscopy [32] to determine protein complex assembly on receptors in live and fixed cell membranes. We studied human embryonic kidney (HEK) cells modified to express monomeric Green Fluorescent Protein (mGFP) labeled hC5aR, exposed to Alexa dye-labeled *S. aureus* toxin components LukS and LukF and imaged using standard total internal reflection fluorescence (TIRF) real time microscopy (Additional file 1: Figure S1a) allowing us to monitor the spatiotemporal dynamics of receptor and toxin molecules in the cell membrane. Our findings indicate that LukS binds on clusters of membrane-integrated hC5aRs. The receptor-bound LukS then binds LukF leading to the formation of a pore that is consistent with previous stoichiometric studies. However, when LukF is bound to the complex, we observe fewer colocalized hC5aRs with toxin in fixed cells, more immobilized toxin complexes in live cells and a significantly reduced Förster resonance energy transfer (FRET) signal, indicating, unexpectedly, that pore formation leads to simultaneous dissociation of the receptors from the complex. In addition, our biochemical data suggests that the dissociated receptor can then be available for additional LukS molecules or the C5a generated during complement activation as a response to LukSF-mediated cell lysis. This new finding suggests that a limited number of receptors can be ‘recycled’ as docking for further toxin. This ensures that a sufficient number of pores will damage nearby phagocytic cells, particularly important when high numbers of C5a anaphylatoxin are blocking LukS, and potentially also enables simultaneous C5a mediated inflammatory response on adjacent cells.

## RESULTS

### Maleimide-labeled LukSF mediates toxicity on human polymorphonuclear (PMN) and HEK cells

To study LukSF pore formation on live cells using single-molecule fluorescence microscopy, single cysteine substitutions on the exposed surface of the cap domain of the individual toxins (Additional file 1: Figure S1b), K288C on LukF and K281C on LukS, were engineered to facilitate maleimide labeling. These were denoted as the modified protein mLukF or mLukS. A second substitution Y113H on LukS was chosen on the stem domain to facilitate pore formation of the LukS mutant (mLukS), based on previous studies [17]. We compared the lytic activity of these mutants to their unmodified wild type equivalents by measuring PMN membrane permeabilization after 30 min toxin exposure using the DNA-binding fluorescent dye DAPI by flow cytometry. DAPI does not penetrate intact cell membranes, and is therefore a good measure for cell permeability and cell death. In this assay each of the wild type toxins was replaced with the modified protein either unlabeled (mLukF or mLukS) or with a single Alexa647 dye molecule label (mLukF* or mLukS*) (Figure 1). All modified toxins induced PMN permeabilisation reaching 100% at ~3 nM (Figure 1a), interchangeable with the wild type equivalents. Only maleimide-labeled mLukF (mLukF*) lost activity and required ~30 nM to reach 100% permeabilization. Since the LukS component mediates the toxin recognition on the target C5aR, we evaluated the binding potency of mLukS and mLukS* on PMNs. In this assay, mLukS was able to inhibit the interaction of FITC-labeled wild type LukS on PMNs equally well as the maleimide-labeled mLukS* (Figure 1b).

**Figure 1.**
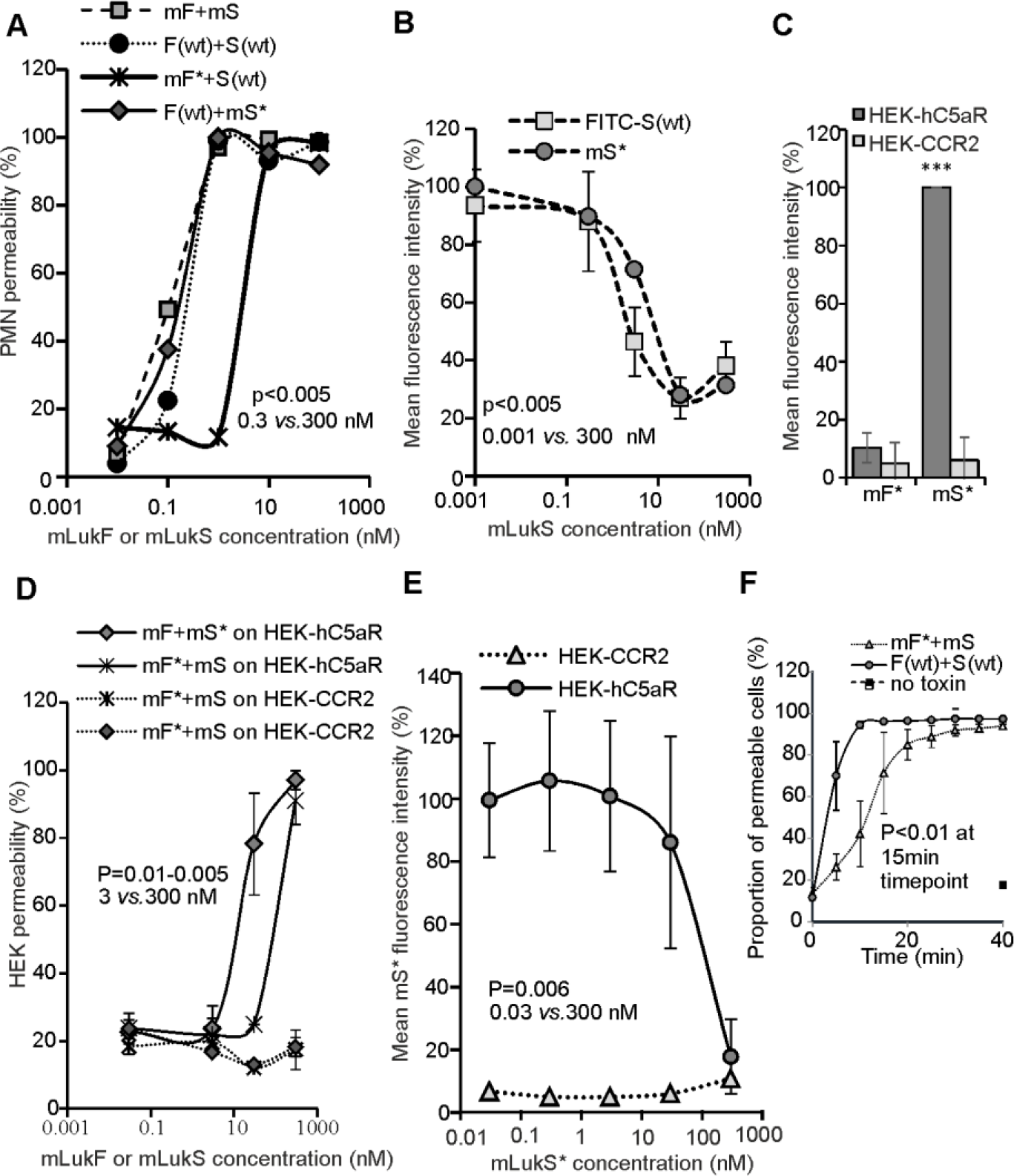
Toxin functionality on PMN and HEK cells. (A) PMN cell permeability in the presence of unlabeled LukSK281CY113H and LukFK288C (mS+mF, number of biological replicates n=2), Alexa-labeled mS* or mF* and wild type LukF (F(wt)) or LukS, (S(wt)) (n=1), compared to PMN cell permeability of S(wt) and F(wt), (n=3); (B) Inhibition of 3 µg/ml of FITC-labeled S(wt) (n=3) and mS* (n=1) binding to PMN cell by mS; Permeability dose dependencies for (A) and (B) are shown with a polynomial spline fit, statistical significance indicated between low (0.3 and 0.001 nM) and high (300 nM) toxin concentrations using Student’s *t*-test. Error bars indicate SD. (C) Column indicating binding responses for mF* on hC5aR cells (n=2). *** Indicates a statistically significant difference (p<0.001) between mS* binding on HEK-hC5aR cells compared to HEK-CCR2 and mF* binding on these cells.(D) Permeability of hC5aR transfected HEK cells using unlabelled mS and mF, and Alexa-labeled mS* and mF* compared to wild type S(wt) and F(wt) (n=2). (E) Inhibition of 3 µg/ml of mS* binding by mS on HEK-hC5aR cells, (n=3). CCR2-transfected HEK cells used as negative controls for toxin binding and lysis in (C, D) n=2 or (C) one representative experiment. Dose dependency shown with polynomial spline fit. Statistical significance calculated between low (0.3 and 0.001 nM) and high (300 nM) toxin concentrations using Student’s *t*-test. (F) Permeability response of hC5aR-transfected HEK cells following incubation with unlabeled mS and Alexa maleimide-labeled mF* or wild type toxins F(wt) and S(wt) (n=3). Statistical significance calculated between 15 min and 0 min time points using Student’s *t*-test. Error bars indicate SD. Percentages of mean fluorescence intensities is shown as relative to the maximum intensity in each individual experiment (B, C and E). Permeability of the cells were analyzed after 30 minutes incubation at +37°C while the inhibition assays were analyzed after 45 minutes incubation at +4°C.

The LukSF toxin is known to be specific towards human cells expressing human C5aR (hC5aR) such as neutrophils, monocytes and macrophages but does not lyse cells that do not express the receptor [18]. To report on the spatiotemporal localization of the receptor and for determining the subunit stoichiometry in any observed receptor clusters we prepared HEK cells expressing hC5aR with a monomeric variant (bearing the obligate-monomer mutation A206K) of green fluorescent protein (mGFP) cloned in the C-terminal end of the receptor. This cell line also forms a monolayer on the coverslip and can be used for introducing a single dye on the cloned receptor, requirements for TIRF and single-molecule imaging. We verified the specificity and activity of the mutated and labeled toxins on the HEK-hC5aR cells. As expected, the toxins lysed only cells expressing hC5aR while control hCCR2 expressing cells, that do not bind LukS [33], remained intact (Figure 1d). We did not observe any binding of mLukF* on the same cells (Figure 1c) which is consistent with previous observations that LukF in the absence of LukS does not interact with PMN [16]. The unlabeled mLukS inhibited binding of mLukS* in a dose-dependent fashion (Figure 1e). HEK cells transfected with hC5aR required higher toxin concentrations for optimal binding and lysis by mLukSF as compared with PMN which is in agreement with previous data for the wild type variants [33]. We did not compare the hC5aR expression levels between these cells as PMNs also express another ligand for LukS, C5LR (or C5aR2) and because C5aR expression levels are not stable in neutrophils but can easily change in natural settings for example as a response to increased C5a levels [34].

To be able to analyze the dynamics of receptor and toxin interactions, we verified the conditions required for HEK cell lysis in time in the presence of mLukF and mLukS. Since the maleimide-labeled mLukF* required higher concentrations for efficient lysis of HEK cells, and because of the loss of molecules during washing cycles, the assay was optimized to have 20-fold excess of mLukF* (600 nM). Following pre-incubation of hC5aR-mGFP expressing HEK cells with LukS(wt) or mLukS, LukF(wt) or mLukF* was added and the cellular uptake of DAPI was measured by flow cytometry every 5 min. The wild type toxins LukF(wt) and LukS(wt) caused >80% cell toxicity within 10 min, while closer to 20 min was required for significant lysis by the mLukF* and mLukS toxin combination (Figure 1f).

### Standard TIRF microscopy of live cells shows LukS colocalizes to hC5aR, and causes cell lysis upon addition of LukF

For life cell imaging we first set the conditions to facilitate data acquisition of dynamic events involved in the formation of LukSF nanopores in hC5aR-mGFP HEK cell membranes. We sampled every 2.5 s at 50 ms exposure time per frame using standard (non single-molecule) total internal reflection fluorescence (TIRF) microscopy, at very low excitation intensity to prevent photobleaching. Cells were first imaged in the absence of toxin. In the green channel we observed mGFP localization consistent with the cell membrane, manifest as relatively high apparent brightness towards the cell boundaries consistent with the cell membrane curving away from the microscope coverslip perpendicular to the TIRF excitation field. Controlled addition of mLukS* (labeled with Alexa647) to the sample Petri dish followed by washing, while imaging simultaneously throughout, resulted in colocalization of the hC5aR and mLukS (Figure 2, Additional file 6: Movie 1), with an image structural similarity index of ~0.75. Further addition of mLukF resulted in complete lysis of the cell, as defined by the observation of explosive release of membrane vesicles, after ~15 min (Figure 2, Additional file 7: Movie 2). Colocalization of hC5aR-GFP, mLukS* (Alexa647) and mLukF* (Alexa594) was also confirmed by three color experiments, imaging cells after addition of toxins and washes until the start of lysis (Figure 2). In the given mLukF concentrations and time frame, where imaging was possible without immediate cell lysis, also free hC5aR could be detected without mLukSF colocalization.

**Figure 2.**
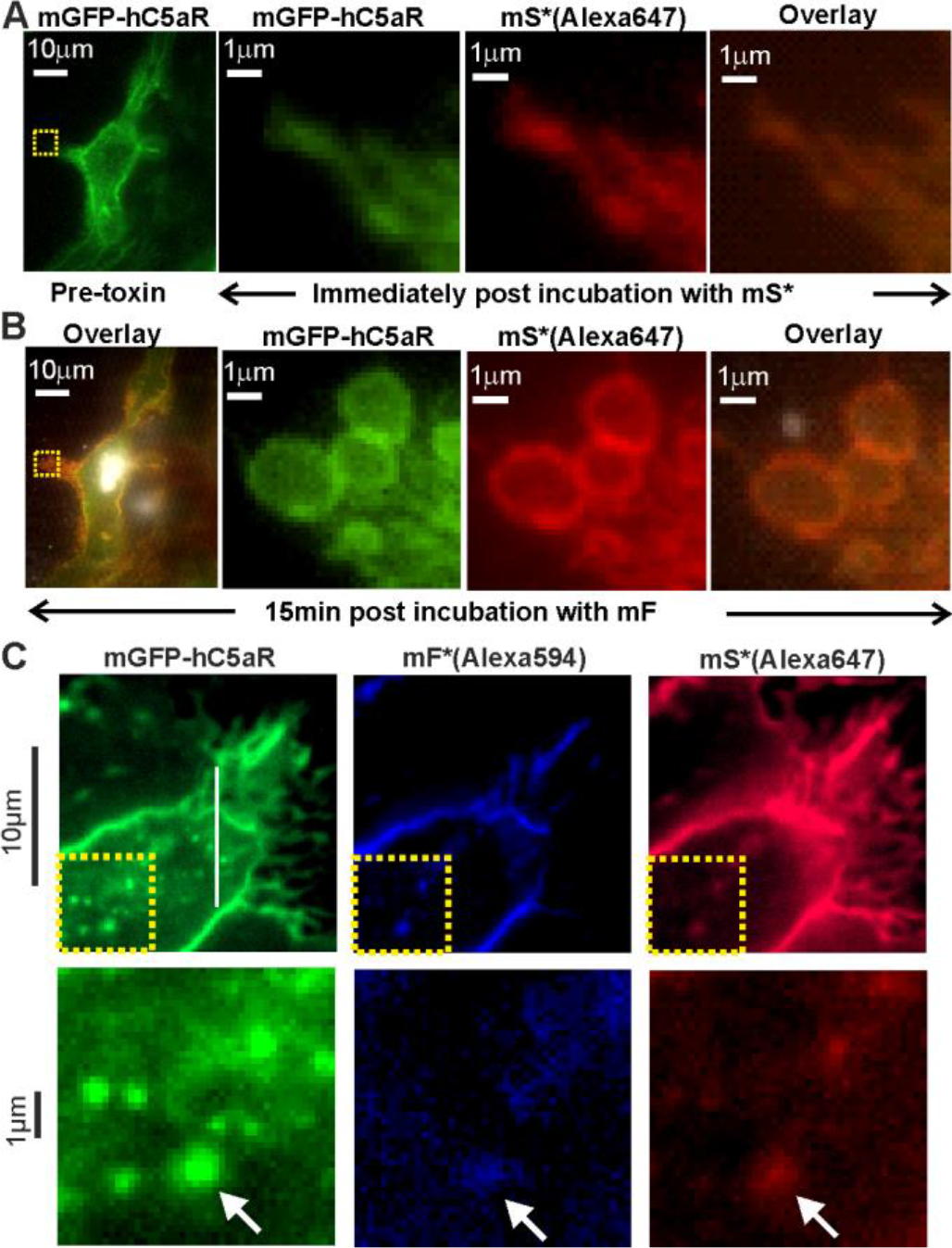
Colocalization of LukS with hC5aR on HEK cells. (A) (left panel) TIRF image of hC5aR-mGFP on the surface of a HEK cell before addition of toxin; (right panels) zoom-in of yellow dashed square of left panel immediately following 2 min incubation with Alexa647-labeled LukSK281CY113H (mS*-Alexa647). (B) Equivalent images of same cell of (B) after > 15 min incubation with LukFK288C (mF). (C) (upper panel) TIRF image of colocalization of Alexa594- and Alexa647-labeled mF and mS (mF*Alexa594 and mS*Alexa647) with hC5aR-mGFP on HEK cells; (lower panel) zoom-in of yellow-dashed square of upper panel with colocalized foci indicated (white arrow).

### Single-molecule TIRF microscopy of live cells shows LukS forms tetramers, and clusters hC5aR before binding LukF

Using higher laser intensity TIRF excitation enabled rapid millisecond single color channel sampling of single fluorophores faster than their molecular mobility in the cell membrane [35], confirmed by imaging antibody-immobilized mGFP and Alexa dyes (Additional file 2: Figure S2). Imaging live hC5aR-mGFP cells in these conditions saturated the camera CCD but after 1-2 min of exposure, photobleaching was sufficient to reduce intensity and allow us to observe several distinct, mobile, circular fluorescent foci at a mean surface density of ~1 per µm^2^ in the planar membrane regions which lie parallel to the TIRF field away from the cell boundaries (Figure 3a, Additional file 8: Movie 3). We monitored the spatiotemporal dynamics of foci in the planar membrane regions using automated tracking software [36] which allowed foci to be tracked for several seconds to a spatial precision of ~40 nm [37], below the diffraction limit, thus enabling super-resolution localization data to be obtained. The measured focus width (defined as the half width at half maximum determined from their pixel intensity profile) was in the range 200-300 nm, consistent with the point spread function (PSF) width of our microscope. By using step-wise photobleaching analysis we estimated stoichiometry values for all detected fluorescent foci by employing a method which quantifies the initial unbleached foci brightness and divides this by the measured brightness for the relevant single dye reporter molecule (Additional file 2: Figure S2) [38]. These foci contained large numbers of receptors with a mean stoichiometry of ~180 (Figure 3b, Table 1). Addition of mLukS and mLukF increased the mean stoichiometry by >50% consistent with the toxin causing receptor clustering.

**Figure 3.**
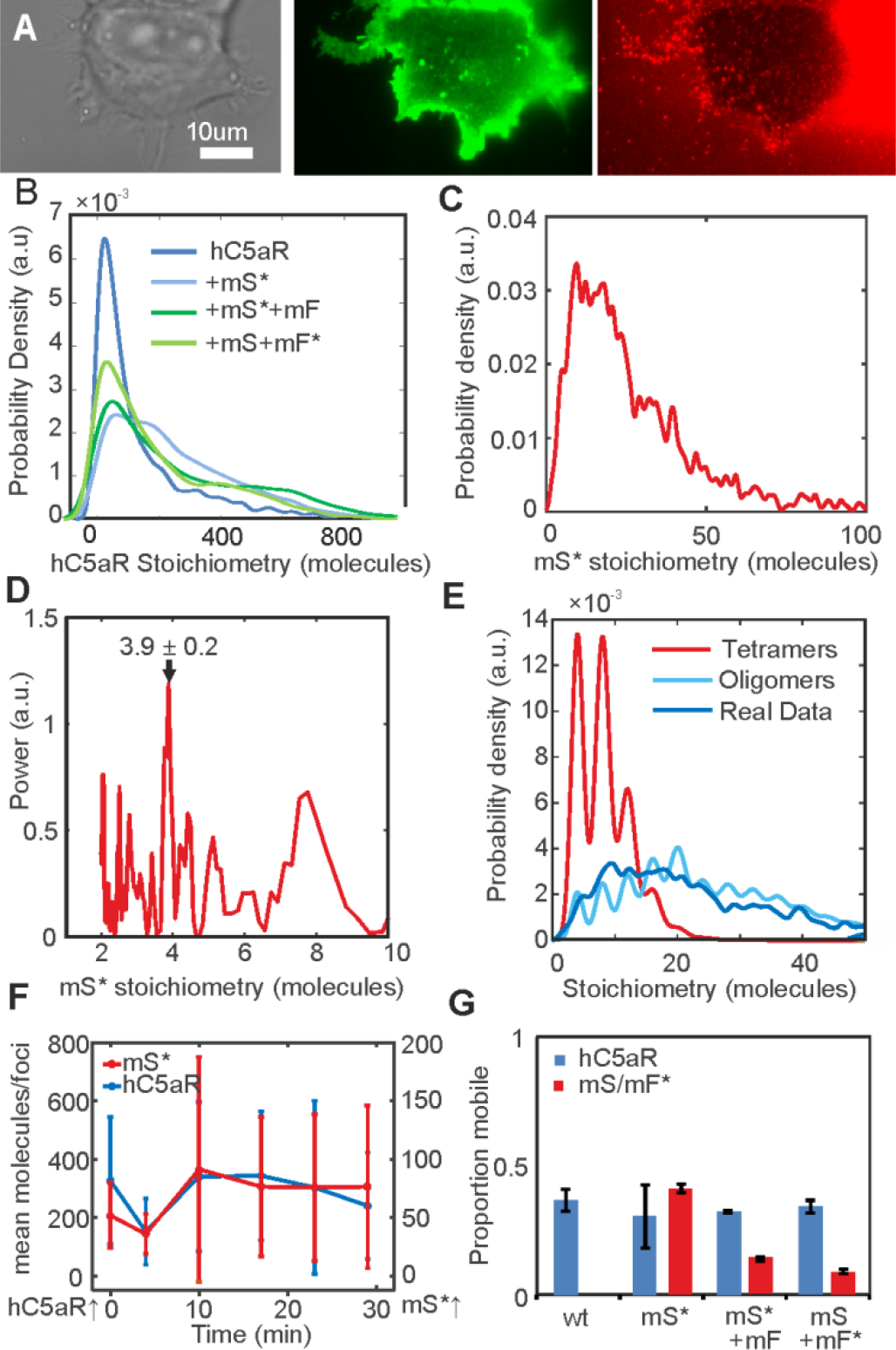
Spatiotemporal dynamics of hC5aR, LukS and LukF in live cells. (A) Images of HEK cells treated with LukSK281CY113H (mS) and Alexa-labeled LukFK288C (mF*) showing brightfield (left), hC5aR-mGFP (middle) and mF* (right). (B) Probability distribution for stoichiometry of hC5aR in absence and presence of Alexa-labeled mS (mS*) and mF*, and (C) of mS* foci, indicating (D) tetramer periodicity from Fourier spectral analysis. (E) A random tetramer overlap model cannot account for mS* experimental stoichiometry data (*R^2^*<0), but a tetramer-multimer model results in excellent agreement (*R^2^*=0.85). (F) hC5aR and mS* stoichiometry as a function of incubation time. Proportion of immobile and mobile colocalized foci in the (G) presence and absence of mS and mF. Error bars show standard error of the mean from n=5-15 image subregions.

**Table 1.** Mean and standard deviation stoichiometry and diffusion constant in live and fixed cells.

|  | Mean [SD]<br>stoichiometry<br>(molecules) | Number of foci | Mobile foci mean<br>[SD] *D*<br>(μ<sup>2</sup>m/s) | Immobile foci mean<br>[SD] *D*<br>(μ<sup>2</sup>m/s) |
| --- | --- | --- | --- | --- |
| C5a | 185[224] | 5346 | 0.48[0.43] | 0.03[0.03] |
| C5a+mLukS* | 281[213] | 8272 | 0.49[0.42] | 0.02[0.03] |
| C5a+mLukS*+mLukF | 291[248] | 6605 | 0.46[0.42] | 0.03[0.03] |
| C5a+mLukS+mLukF* | 223[202] | 4981 | 0.47[0.40] | 0.02[0.03] |
| mLukS* | 29[22] | 999 | 0.44[0.45] | 0.03[0.03] |
| mLukS*+mLukF | 84[89] | 841 | 0.34[0.35] | 0.01[0.02] |
| mLukS+mLukF* | 6[4] | 557 | 0.40[0.44] | 0.02[0.03] |

Fixed cells:
|  | Number and (%) of<br>colocalized C5a foci | Number and (%) of<br>C5a foci not<br>colocalized | Mean [SD]<br>stoichiometry of<br>colocalized C5a foci<br>(molecules) | Mean [SD]<br>stoichiometry of C5a<br>foci which are not<br>colocalized<br>(molecules) |
| --- | --- | --- | --- | --- |
| C5a+mLukS* | 89(32) | 193(68) | 36[26] | 35[24] |
| C5a+mLukS*+mLukF | 84(7) | 1079(93) | 37[25] | 32[24] |
| C5a+mLukS+mLukF* | 59(83) | 283(17) | 26[20] | 26[20] |
|  | Number and (%) of<br>colocalized Luk<br>foci | Number and (%) of<br>Luk foci not<br>colocalized | Mean [SD]<br>stoichiometry of<br>colocalized Luk foci<br>(molecules) | Mean [SD]<br>stoichiometry of Luk<br>foci which are not<br>colocalized<br>(molecules) |
| mLukS* | 88(7) | 1198(93) | 72[47] | 61[44] |
| mLukS*+mLukF | 86(88) | 658(12) | 121[95] | 136[94] |
| mLukS+mLukF* | 60(5) | 1186(95) | 20[16] | 18[15] |

Imaging mLukS* incubated with hC5aR-mGFP cells revealed distinct foci (Figure 3a, Additional file 9: Movie 4). The probability distribution of mLukS* stoichiometry values in live cells in the absence of mLukF is shown in Figure 3c, rendered using a kernel density estimation which generates an objective distribution that does not depend upon the size and location of subjective histogram bins [39]. We measured a broad range of stoichiometry values, spanning a range from only a few LukS molecules per foci to several tens of molecules, with a mean of ~30 molecules per foci. Closer inspection of the stoichiometry values indicated an underlying periodicity to their probability distribution, which we investigated using Fourier spectral analysis [40]. The resulting power spectrum (Figure 3d) indicated a fundamental peak equivalent to a stoichiometry of 3.9 ± 0.2 molecules, suggesting that foci are composed of multiples of tetrameric mLukS* complexes.

Fluorescent foci, if separated by less than the diffraction-limited PSF width of our microscope, are detected as a single particle but with higher apparent stoichiometry. We therefore tested the hypothesis that the observed mLukS* foci stoichiometry distribution could be explained by the random overlap of isolated mLukS* tetramer foci. To do so we modeled the nearest-neighbor separations of individual mLukS* tetramers in the cell membrane as a random Poisson distribution [41] and used sensible ranges of tetramer surface density based on our single particle tracking results (Additional file 3: Figure S3). However, all random tetramer overlap models we explored showed poor agreement to the observed experimental stoichiometry distribution, but we found that random overlap of multimers of tetramers could account for the stoichiometry distribution well (Figure 3e). Optimized fits indicated that the random overlap of mLukS* foci with a stoichiometry in the range 4-20 molecules were able to best account for the experimental data. As hC5aR is clustered, this likely accounts for the clustering of mLukS* but not its tetrameric periodicity. These results are consistent with mLukS* binding to clusters of hC5aR as tetramers or forming tetrameric sub-structures.

We tested if there was a dependence of foci stoichiometry on incubation time with leukocidin. Acquiring a time course for mLukF* accumulation following pre-incubation of cells with mLukS was not feasible since unbound mLukF* had to be washed from the sample to prevent a prohibitively high fluorescent background. However, we were able to acquire time courses in which mLukF was added to cells that had been pre-incubated with mLukS*. For these, the mLukS* foci stoichiometry distribution was measured as a function of time after mLukF addition for several different fields of view, each containing typically ~5 cells. We found that the mean hC5aR foci stoichiometry indicated no obvious correlation to mLukF incubation time (Figure 3f), however the mean mLukS* foci stoichiometry increased with time (p <0.05).

By calculating the mean square displacement (MSD) as a function of time interval (τ) for each tracked foci we could determine its apparent microscopic diffusion coefficient (*D*). The distribution of *D* for hC5aR and mLukS*/mLukF (Additional file 4: Figure S4) had similar low value peaks at ~0.05 µm^2^/s, consistent with immobile foci tracked with our localization precision spatial precision of 40 nm. Several mobile foci were also seen, which diffused at rates up to ~5 µm^2^/s. Based on the measured width of the immobile peak width on these distributions we set a threshold of 0.12 µm^2^/s to categorize foci as either immobile, which indicated a mean *D*=0.025 ± 0.030 µm^2^/s (±SD), or mobile, which indicated a mean *D*=0.47 ± 0.40 µm^2^/s (Table 1). Plots of the measured MSD *vs*. τ relations for mobile foci indicated a linear dependence indicative of free Brownian (i.e. ‘normal’) diffusion. However, similar plots for immobile foci indicated an asymptotic dependence consistent with confined diffusion [42], whose plateau was equivalent to a confinement diameter of ~400 nm (Additional file 4: Figure S4). The relative proportion of mobile foci was ~35% of tracked foci for hC5aR, regardless of toxin and similar for mLukS in the absence of mLukF. Addition of mLukF, caused a drop in the mobile proportion by a factor of ~3 (Figure 3g) suggesting that LukF causes insertion of the complex and possible disassociation of the LukSF complex from the hC5aR.

### Single-molecule TIRF microscopy combined with colocalization analysis of fixed cells suggests LukSF dissociates from the receptor

Due to the high image frame rate of single-molecule TIRF microscopy, we were not able to simultaneously image two color channels on our microscope, rather each channel was imaged separately in the same cells. Therefore to determine whether the toxin remains bound to the receptor, and to quantify the relative stoichiometry of components, we imaged fixed cells, halting cell lysis, using the same two spectrally distinct green/red dyes of mGFP and Alexa647 to label receptor and toxin components, respectively, as for the live cell experiments. We imaged cells incubated with mLukS*, followed by incubation with mLukF (Figure 4a upper panels) as well as simultaneously with mLukS+mLukF* (Figure 4a lower panels) and observed foci with similar stoichiometries (Table 1) to live cells but colocalized with hC5aR. Around 32% of the hC5aR foci were found colocalized in the presence of mLukS* dropping to <10% in the presence of mLukF (Figure 4b). This low percentage was within our estimate of the degree of random colocalization between the green and red fluorophores, entirely down to chance, of ~10%. This suggests that in the presence of mLukF, the toxin is not colocalized with the receptor and that mLukF causes disassociation from hC5aR. The stoichiometry values for detected green hC5aR-mGFP foci were calculated and plotted against the equivalent stoichiometry estimates for colocalized red foci of mLukS* and mLukF* respectively (Figue 4c and Additional file 5: Figure S5). In the presence of mLukS* but in the absence of mLukF, the hC5aR-mGFP foci stoichiometry showed an approximately linear dependence on number of associated mLukS* molecules, suggesting that each colocalized mLukS* molecule was associated on average with ~4-5 hC5aR molecules. In the presence of labeled or unlabeled mLukF no dependence was observed (Additional file 5: Figure S5, R^2^<0) consistent with random association between toxin and receptor. These results are unlikely to be due to fluorescence quenching, as it would need to be near 100% quenching to detect no Alexa fluorescence in the hC5aR-mGFP foci and the drop in colocalization is observed independent of the labeled toxin used, either mLukS* with mLukF or mLukF* with mLukS.

**Figure 4.**
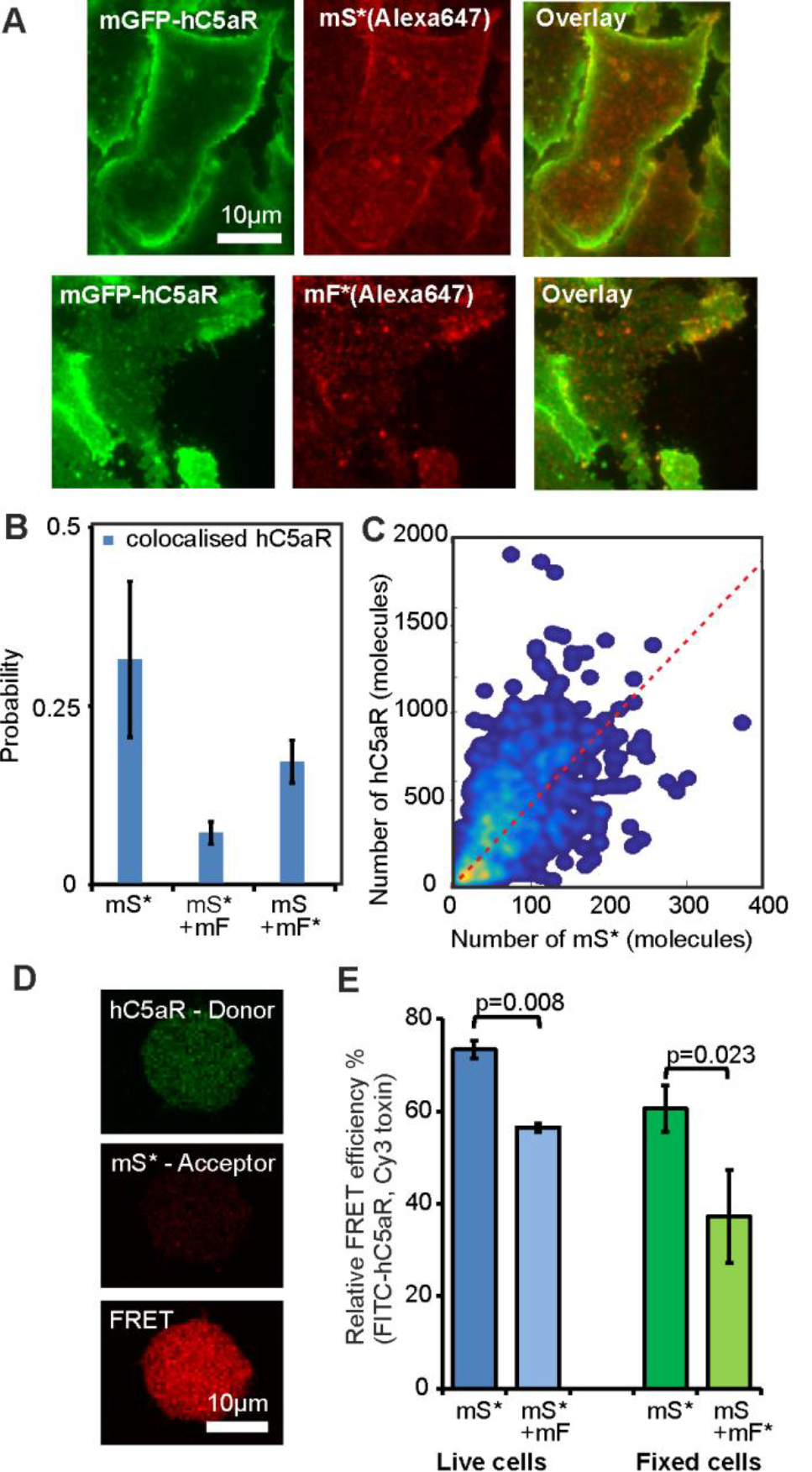
Relative stoichiometry of hC5aR, LukS and LukF in fixed cells. (A) Micrographs of fixed hC5aR-mGFP HEK cells treated with LukSK281CY113H (mS) and LukFK288C (mF) showing hC5aR-mGFP (left) and Alexa647 (middle) and merge (right) on Alexa-labeled mS (mS*) above and mF (mF*) below (B) Proportion of foci colocalized and not colocalized, treated with mS, mS*+mF and mS+mF* for hC5aR. Error bars show standard error of the mean from n=4 image subregions. (C) Heatmap of correlation between hC5aR and mS stoichiometry (red dash line indicates 4 mS per hC5aR molecule), R^2^ ~0.15 (D) and (E) FRET images and efficiencies. The FRET experiment was performed in live and fixed sortase-tagged FITC-hC5aR expressing cells. Live cells (number of biological replicates n=2) were incubated in the presence of Cy3-labeled mS* for 1 h at +4°C and washed, after which unlabeled mF was added. FRET was analyzed before (mS*) or after addition of mF (mS*+mF). FRET from fixed cells (n=3) was analyzed in the presence of mS* or unlabeled mS and Cy3-labeled mF* respectively (n=2). Statistical significance between cells with only mS and both of the toxin components, mS and mF, was analyzed using Student’s *t*-test. Error bars indicate SD.

### Live whole cell FRET and biochemical measurements also support LukSF disassociation

We performed FRET experiments on FITC sortase-labeled hC5aR and Cy3-labeled mLukS or mLukF, as donor and acceptor respectively, in live cells to further probe the association between toxin and receptor. A FRET signal from whole cells of 75% efficiency was observed, with a statistically siginfcant drop (p=0.008, Student’s *t*-test) to 56% when incubated with unlabeled mLukF (Figure 4d and e), as would be expected if the complex formation leads to dissociation of the toxin from the receptor. In order to examine possible FRET between hC5aR and Cy3-labeled mLukF we performed similar experiments on fixed cells. In these experiments a FRET efficiency of 60% was observed between hC5aR and labeled mLukS dropping below 40% (p=0.023, Student’s *t*-test) between hC5aR and labeled mLukF. As expected, no FRET signal was observed in the negative control where only Cy3-labeled mLukF was present. These results are also consistent with the finding that hC5aR dissociates from the LukSF pore, although conformational or local environment changes cannot be ruled out with FRET alone since the relatively high remaining signal might in principle also indicate remaining association or other inter or intra hC5aR-Luk interaction. The greater drop in FRET when measured with mLukF compared to mLukS might be caused by the 3-4 nm further distance of LukF from hC5aR.

To further confirm that the LukSF complex dissociates from the target receptor we used a monoclonal PE-labeled anti-CD88 antibody to detect the liberation of free hC5aR receptors on the cell membrane upon LukSF formation. We first confirmed the ability of both C5a and wild type LukS to compete for binding of the anti-CD88 antibody to the hC5aR expressing HEK cells. Both ligands showed clear inhibition of anti-CD88 binding at 100 nM concentrations while LukF was ineffective (Figure 5a). However, when the hC5aR expressing cells were incubated with 100 nM of LukS followed by incubation with increasing concentrations of wild type LukF to form an active toxin, a statistically significant increase in anti-CD88 binding was detected at a LukF concentration of 1nM when compared to no LukF (0 nM). Addition of a control protein Ecb did not change anti-CD88 binding. Cell permeabilization was measured in parallel and proved to be “sublytic” enabling proper detection of liberated anti-CD88 without significant cell lysis (% of lysed cells < 10%) (Figure 5b). At a 3 nM LukF concentration the proportion of dead cells increased above 10% which determined the maximum concentration and increase in anti-CD88 binding that could be measured. The changes in C5aR mobility, colocalization, and FRET with addition of LukF, combined with the biochemical evidence of anti-CD88 rebinding on hC5aR upon LukF(wt) addition on LukS(wt) coated cells are strongly indicative of disassociation of the LukSF complex.

**Figure 5.**
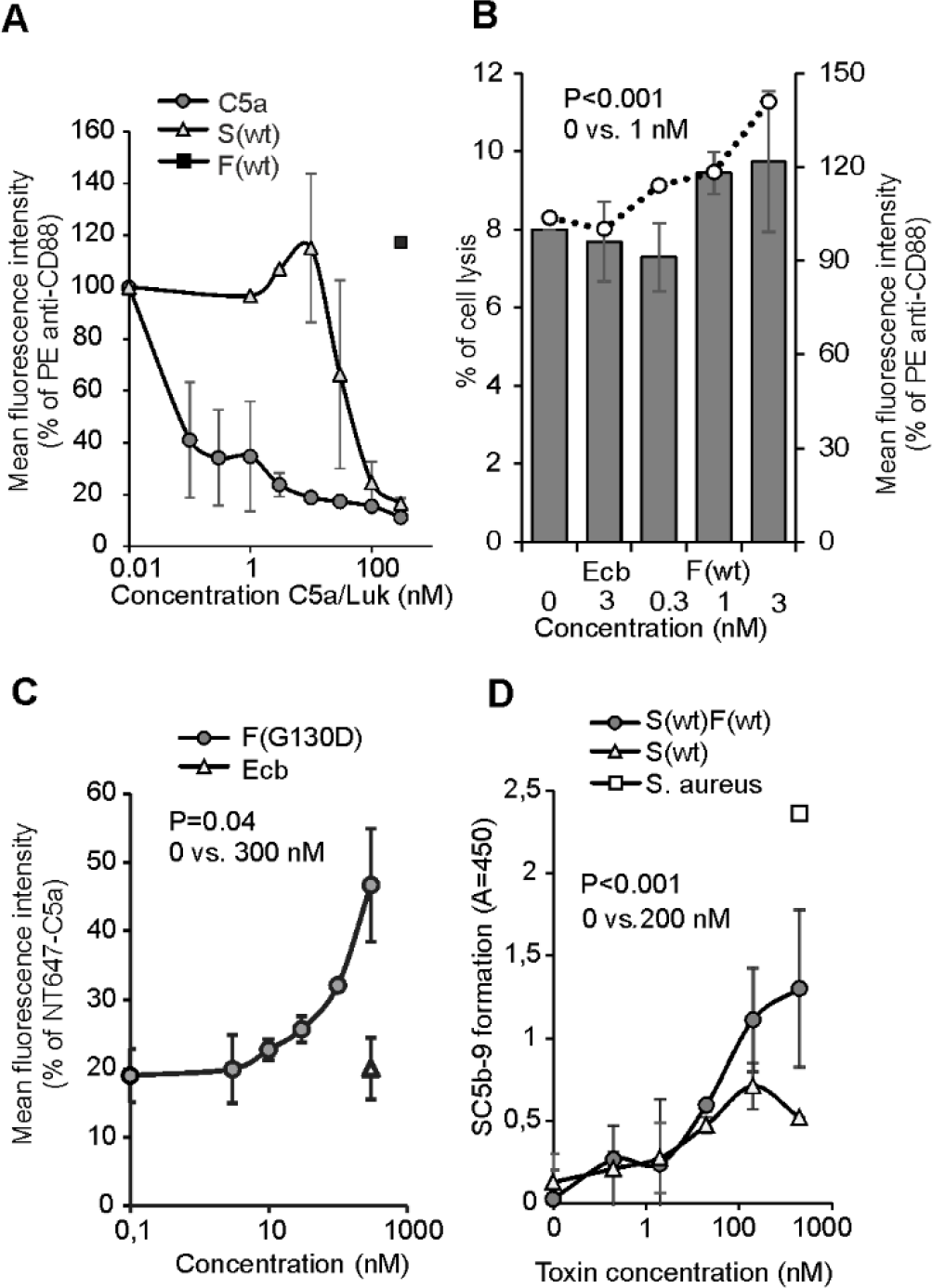
LukSF dissociation and rebinding of C5a on hC5aR expressing cells. (A) Inhibition of anti-CD88 binding on hC5aR expressing HEK cells using increasing concentrations (horizontal axis) of wild type LukS, S(wt) and C5a. Wild type LukF, F(wt), is used as a negative control for inhibition of anti-CD88 binding (number of biological repeats n=2). The values are normalized against the maximum binding observed with only anti-CD88. (B) Disengagement of hC5aR from LukSF was observed as increase in PE conjugated anti-CD88 binding (right vertical axis, indicated with bars) on S(wt) pre-coated cells using increasing but sublytic concentrations (horizontal axis, indicated with dots) of F(wt). The values are normalized against the maximum binding observed with LukS incubated cells with only anti-CD88. Minimal cell lysis (% of lysed cells, left vertical axis) detected in F(wt) concentrations below 3 nM (n=3). (C) Rebinding of constant amount of NT647 labeled C5a (647-C5a) on hC5aR upon LukSF formation analyzed by incubating S(wt) pre-coated cells with increasing concentrations (horizontal axis) of LukF mutant G130D that associates with LukS but does not lead to cell lysis (n=2). The values are normalized against the maximum binding observed with only 657-C5a. (D) Effect of LukSF mediated cell lysis on complement activation and C5a formation on full blood measured by using C5b-9 as a marker for complement activation in plasma (n=3). Maximal C5a formation is observed by incubating full blood with live *S. aureus* bacteria. Ecb (B and C), F(wt) (A) or S(wt) (D) are used as negative controls in the assays. Percentages of mean fluorescence intensities is shown as relative to the maximal intensity in each individual experiment (A-C). Statistical significances are calculated using Student’s *t*-test. Error bars indicate SD.

### Dissociation of LukSF pores from hC5aR allows rebinding of C5a or intact LukS on the receptor

Since C5a is the natural ligand for hC5aR and can outcompete binding of LukS on the receptor we next analyzed whether LukSF formation and disengagement of hC5aR would allow rebinding of C5a on the receptor. We specifically chose to analyze binding of labeled C5a and not LukS in the presence of increasing concentrations of LukF. This is because adding labeled LukS to cells coated with LukS (and then washing) together with LukF will give several possibilities for association: for example, new pores for free or unoccupied C5aR; intercalation with present bound LukS and lukF in pores; binding to free or unoccupied C5aR but without pore formation. To detect C5a rebinding at higher LukF concentrations we used a G130D LukF mutant that interacts with LukS but does not cause cell lysis. Non lytic activity of this mutant in this assay was confirmed by DAPI staining that showed minimal cell lysis even at higher concentrations (9% at a 300 nM LukFG130D concentration). At a 300 nM concentration a significant increase in C5a binding was detected indicating that LukSF dissociates from hC5aR enabling simultaneous rebinding of an hC5aR interacting ligand (Figure 5c). On the contrary, addition of the control molecule, Ecb, did not cause increase in C5a binding suggesting that this was due to rebinding of C5a to disengaged hC5aR and not for example because of increase in receptor expression. This assay showed that C5a can potentially interact with these cells that are attacked by LukSF. Because C5a is a potent anaphylatoxin that is generated during complement activation and potentially plays a crucial role in *S. aureus* infections [43] we next analyzed whether LukSF could lead to complement activation and C5a formation in an *ex vivo* full blood assay. We used soluble C5b-9 as a marker for terminal complement activation and C5a formation, not C5a, because LukS is known to compete with C5a for binding to hC5aR on neutrophils [18]. The presence of 200 nM of LukSF clearly increased formation of soluble C5b-9 compared to full blood without any toxin or only LukS (Figure 5d), indicating that LukSF mediated cell lysis increases C5a formation and potentially also inflammation in the site of infection.

## DISCUSSION

To determine the stoichiometry of the toxin components without immobilizing the protein on a surface or within a crystal we implemented our fluorescence imaging method which allows us to monitor the actual pore formation mechanism within a living cell, including the target receptor crucial for the complex formation. This kind of study on protein complex formation has not been done before due primarily to the difficulty of labeling the components and the high native fluorescence background in mammalian cells. Our covalent labeling strategy and high excitation intensity TIRF microscopy, combined with advanced image analysis tools, opens the way for further studies into many other pore forming toxins and processes involving membrane bound protein complex formation.

The finding that the toxin complexes are found in receptor clusters indicates that lysis of cells depends on the local density of close proximity hC5a receptors that will initiate the pore formation process by docking LukS close to the cell membrane such that four hC5aR-LukS dimers (assuming that one LukS binds only one hC5aR, although hC5aR are randomly clustered on the membrane) can interact with the free non-bound LukF that will eventually form an octamer (i.e. 4 by 4) and a functional pore with LukS. These data also suggest that when proper assembly in an octamer is ongoing/complete, hC5aR will dissociate from the complex and, at the same time, can interact with the C5a that is formed during complement activation amplified by LukSF-mediated cell lysis itself. This is logical because, in addition to invading microbes, apoptotic and necrotic cells are known to activate complement [44]. The triggering of local complement activation by toxin damaged cells and the release of locally generated C5a (Figure 5) and its interaction with adjacent cells such as endothelial or lung epithelial cells [45] could explain the mechanism behind the exacerbated inflammation characteristics exhibited in necrotizing pneumonia. This is an important finding, suggesting that the cause of infection can dramatically affect the magnitude of the inflammatory response and is highly dependent on the dynamics of microbial molecules interacting with human receptors. In addition, the disengaged hC5aR is possibly also available for new toxins to bind, thus allowing the receptor to be recycled and reused by additional LukS molecules. Our finding that C5a can rebind, doesn’t only suggest a mechanism for exaggerated immune response by LukSF, but also indicates that the dissociated free hC5aR does not change in conformation due to previous contact with LukS and therefore would also be available for binding with LukS. We find that roughly half of LukS complexes are immobile prior to LukF binding, but that LukF binding then results in mostly immobile LukSF complexes. Speculatively, this result may suggest that LukS binds hC5aR initially and then inserts itself transiently in the membrane phospholipid bilayer via the exposed hydrophobic residues, following binding of LukF molecules to LukS. This stable insertion of the LukSF complex into the cell membrane then leads to pore formation across the whole cell membrane.

To characterize the hC5aR interaction with LukSF at a molecular level, we used malemide-labeled toxins and HEK cells that expressed only hC5aR and not the second docking target for LukSF, which are both present on human PMNs [46]. We verified that the interaction between maleimide-labeled toxin component LukS and the cell surface receptor is required for the target recognition and cell lysis similarly as shown before for wild type LukS [18] both for human PMNs and hC5aR expressing HEK cells that were chosen for TIRF imaging because of their stability and ability to form monolayers on the microscopy cover slip.

We observe clusters of pre-established hC5aR in the absence of LukS or LukF, but the addition of LukS or LukF significantly increased the mean cluster stoichiometry. Fourier spectral analysis combined with foci overlap modeling suggests that LukS complexes comprise a multimer of 4-5 hC5aR subunits each of which contain 4 LukS molecules, even before addition of LukF. Our findings are consistent with the hetero-octamer model of 4-plus-4 LukS/LukF subunits [17, 30, 31], however, our colocalization analysis also indicates the presence of stable complexes of LukS with hC5aR independent of LukF.

By characterizing the mobility of hC5aR and LukS in live cells we find that roughly half of hC5aR and LukS foci diffuse relatively freely in the cell membrane while the remainder are confined to zones in the membrane of ~400 nm effective diameter. However, when LukF is present > 90% of LukS foci become immobile (confined). If LukS were to undergo a conformational change following LukF binding then this may potentially expose hydrophobic residues that could fascilitate insertion of the toxin into the hydrophobic interior of the phospholipid bilayer. This hypothesis is strongly supported by the β-barrel prepore-pore formation putative mechanism of γ-hemolysin. Here the residues responsible for binding with the phospholipid head group are located at the bottom of the rim domain whereas the stem domain forms an antiparallel β-barrel of which the bottom half comprises the transmembrane portion of the pore [31]. This change from receptor associated LukS to cell membrane associated LukSF complex can be seen as a change in the proportion of mobile (receptor associated LukS) and immobile (toxin complexes inserted into cell membrane) foci detected in live cells. GPCRs similarly are known to have heterogeneous mobility and lateral distribution properties in living cells at different states for example before and after activation [47].

Crystallographic evidence from the monomeric LukF and LukS components and the intact γ-hemolysin pore suggests that the pore is octameric formed from 4-plus-4 LukF/LukS subunits [30, 48, 49]. Our findings support this octamer model but unlike previous studies also indicate that LukS pre-forms into a tetramer without LukF and that formation of this tetramer is facilitated by close proximity C5aR clusters. The presence of LukS tetramers in the absence of LukF cannot be further explained by our data. It is, however, possible that if LukS molecules would be associated as a tetramer when bound on the receptor the conformational changes on LukS caused by interactions with LukF should enable association of the LukF subunits to the complex. According the data presented here this is possible because in these assays we first enabled LukS to bind on the receptors and eliminated the effect of freshly formed complexes by free unbound LukS by a washing step before addition of LukF. Each octamer component consists of cap, rim and stem domains. Here, the cap domain contains the site for LukS/LukF interaction while the stem domain unfolds and forms the transmembrane β-barrel upon pore formation. Within crystallization the 2-methyl-2,4-pentanediol (MPD) molecules are bound at the base of the rim domain, and recognized by Trp177 and Arg198 residues, that may participate in recognition of the phospholipid bilayer as suggested in a crystal structure of the LukF monomer [50]. In contrast, the structure of the γ-hemolysin suggests a membrane interaction site within residues Tyr117, Phe119 and Phe139 on the same toxin component [30]. The crystal structure of LukED determined recently reveals important details of the residues on LukE required for receptor identification [51]. This component corresponds to the receptor binding component LukS on the LukSF complex, scanning mutagenesis indicting that LukS residues Arg73, Tyr184, Thr244, His245 and Tyr250, and to a lesser extent Tyr181, Arg242 and Tyr246, are involved in binding to the neutrophil surface [52].

These results suggest that further binding sites for hC5aR on LukS could be possible in addition to those identified in the LukS rim domain [52]. However, since the binding of LukS to neutrophils is inhibited by the C5a it is likely that LukS has only one binding site on the receptor [20]. This is also supported by the similar inhibition profiles of LukS and C5a towards anti-CD88 binding on hC5aR shown in this study. Therefore, the association of LukS with approximately 4-5 hC5aR molecules could be explained by the previous suggestion that C5aR forms homo-oligomers in living cells [53]. Our findings imply that LukSF assembly is dependent on hC5aR cell membrane area density as opposed to the effective hC5aR concentration when calculated over the whole of a target cell volume, such that even when hC5aR cellular expression levels are low, for example when inflammatory mediators are formed to limit the inflammation [34], a cell lysis response may still be achieved through the efficient targeting of receptor clusters and recycling of the receptor molecules in the cell membrane to be re-used by free non-bound LukS to get engaged in octamer pore formation. It is possible that overexpression of hC5aR on HEK cells could lead to an increased ability to form hC5aR foci with many receptors especially because of intracellular GFP tag although we mitigated this possibility by cloning the receptor with a monomeric variant of GFP that does not cause GFP dimerization. Other studies, however, have shown that hC5aR forms clusters of homodimers or heterodimers with the second C5a receptor C5LR (C5aR2) or other GPCRs like CCR5 especially under high concentrations of C5a [53-55].

Previous *in vitro* studies on LukSF pores formed on human leukocytes and rabbit erythrocytes have found evidence for both octamers and hexamers, but importantly both suggest a LukS/LukF ratio of 1:1 [17, 56, 57]. Interestingly we did not observe any correlation to the number of hC5aR present with LukF incubation time once LukF was already bound to LukS. Moreover, when LukS was incubated with LukF using sortase-labeled hC5aR cells a significant reduction was observed in the FRET efficiency signal between LukS and C5aR. It is unlikely that the reduction that was observed in FRET efficiency would be due to a conformational change because the cysteine mutation used for maleimide labeling was designed to be exposed on the cap domain of LukS and LukF (Additional file 1: Figure S1) that in light of the structural data undergoes minimal or no conformational changes during complex assembly [31]. Since our biochemical assays indicate that LukF does not bind directly to hC5aR expressing cells and, that binding of LukF to LukS results in an increased distance between the receptor and the complex, this suggests that LukF binding to LukS results in LukS dissociating from the receptor, released as a newly formed LukSF complex.

We cannot directly determine the cause of this behavior in our present study, however one explanation may lie in the conformational change during the prepore-to-pore transition that has been shown to occur on γ-hemolysin complexes subsequently after binding of LukF to LukS [30, 31]. Interestingly, this same study shows that during the pre-pore state the space for the transmembrane region is occupied by the rim domain of the adjacent octamer in a LukSF crystal. One explanation for these observations, that remains to be explored, is that in addition to the stem domain the residues within the rim domain that interact with the receptor might also have different orientations in the pre-pore state when compared to the pore state. In addition to using the maleimide labeled mLukF and mLukS and fluorescence microscopy the putative dissociation of the hC5aR from the LukSF complex was further verified by using the wild type LukS and LukF proteins in an assay where LukS-coated hC5aR cells were incubated with increasing concentrations of LukF. Here, increase in anti-CD88 binding also clearly indicates LukSF dissociation. In all of the assays where we could observe 20-30% receptor dissociation we used sublytic concentrations of LukF to be able to measure healthy cells with normal membrane fluidity and natural behavior rather than dead cells. This kind of receptor disengagement has been shown before by at least the cytotoxin intermedilysin which interacts with a GPI-anchored complement regulatory molecule on the cell membrane [58]. Moreover, dissociation of LukSF complex is also supported by electron microscopy of LukSF on human leukocyte membrane fragments. Here, the ring-shaped oligomers with outer and inner diameters of 9 nm and 3 nm were shown without a receptor [57].

In this study we also show, for the first time to our knowledge, that the dissociated receptor can be reused by free unbound C5a. This indicates that LukS binding on the receptor does not change receptor conformation and thereby can putatively also be reused by additional LukS. In our full blood model we observed that LukSF mediated cell lysis clearly increased complement activation and C5a formation. The increase in C5a concentration in the site of infection could potentially limit the availability of hC5aR for LukS molecules on neutrophils and thereby reduce lytic activity of the toxin as C5a has previously been shown to reduce LukSF mediated lysis *in vitro* [18]. Rebinding of C5a on the receptor may therefore indicate that in natural settings where all components (i.e. LukS, LukF and C5a) are present C5a can outcompete binding of LukSF on the target cells. Therefore, recycling of the receptor could be one strategy for the toxin to ensure that a sufficient number of pores will damage the cells especially when limited number of receptors are available.

There are several steps on the leukocidin complex assembly that may be critical for the function of the toxin. Based on our observations we provide new information on leukocidin-receptor interactions and propose two additional stages to the processes of pore formation and the mechanism by which LukSF potentially induces inflammation (Figure 6a). Stage 1 is the binding of LukS to hC5aR clusters. The first step in this process is the target recognition of LukS binding to the membrane receptor. We detected LukS-hC5aR complexes on clusters of receptors indicating that pore formation takes place in these clusters. Stage 2 is the binding of 4 LukF molecules to to 4 LukS molecules resulting in a hetero-octamer LukSF nanopore in the neutrophil cell membrane. Stage 3 is then the dissociation of the receptors from the LukSF complex enabling the repcetor to be resued for subsequent binding free unbound ligand to generate more nanopores in the cell membrane and enhance the damage to the neutrophil. We measured a correlation between the number of LukS and hC5aR molecules present in LukS-hC5aR complexes, but with no obvious correlation between the number of LukF and hC5aR molecules when LukF was added to the LukS-hC5aR complex. In addition to previous studies [14] we suggest that LukSF-mediated cell lysis and dissociation from hC5aR can potentially amplify *S. aureus* mediated inflammation in the site of infection (Figure 6). The direct lysis of neutrophils is enhanced by newly formed LukSF complexes that are formed on the cell membrane hC5aR via reattachment of new LukS. Neutrophil lysis activates the complement system and the newly generated C5a induces cytokine/chemokine production and neutrophil chemotaxis via the C5a/C5aR signaling pathway on adjacent cells. Furthermore, the increased vasodilation and vascular permeability (Figure 6b) leads to massive neutrophil accumulation and tissue injury at the site of bacterial infection [59].

**Figure 6.**
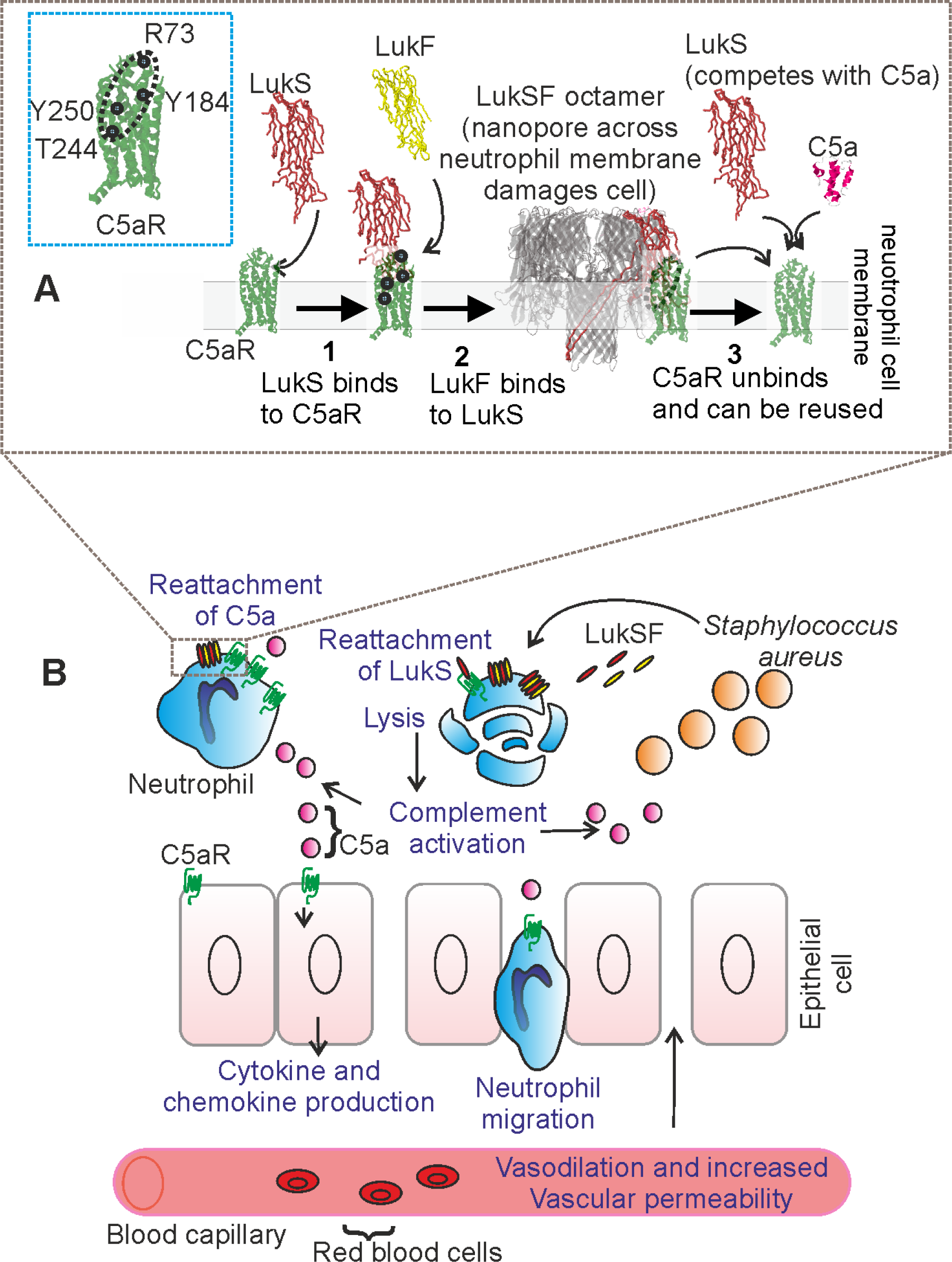
Model for LukSF-receptor binding and the mechanism of LukSF-induced inflammation. (A). LukS (PDB ID: 1T5R) binds on hC5aR (structure based on angiotensin receptor data PDB ID: 4YAY, cyan dashed box) as a soluble monomer on the cell membrane. Each LukS monomer binds one hC5aR molecule via the receptor interacting residues R73, Y184, Y250, T244 (marked with blue dots) within a cluster of approximately 4-5 hC5aR homo-oligomers Upon binding to hC5aR LukS exposes residues for LukF (PDB ID: 1LKF) binding (interface indiacted by dashed ellipse). In these tight clusters each LukF can bind to two LukS monomers via two interfaces. Binding of LukF on LukS and formation of the octameric pore (PDB ID: 3B07) causes dissociation of the receptors from the complex because of leakage of the cell membrane and possibly also since the receptor binding region (marked with a circle) is buried between the monomers in the complex. The detached hC5aR molecule can be reused by its ligands LukS or C5a anaphylatoxin (PDB ID: 1KJS).(B) Zoom out of (A), illustrating the putative mechanism of LukSF induced inflammation.

## CONCLUSIONS

In summary, our findings that the receptors of targeted host cells dissociate rapidly from the leukocidin complex upon formation of a harmful toxin pore, freeing up mobile receptor ‘seeds’ that can diffuse to other parts of the cell membrane, suggest a hitherto undiscovered strategy used by microbes to kill human immune cells. This enables a limited number of receptors to be recycled as docking for the leukocidin or potentially the anaphylatoxin C5a to ensure that enough pores will form to damage the host cell and simultaneously maintain or possibly amplify the inflammation in the site of infection. This discovery may generalize to other bi-component toxins which employ a similar docking receptor like the C5aR receptor, including the family of *Staphylococcal* bi-component leukocidins of HlgC/HlgB, HlgA/HlgB, LukE/LukD (CXCR1, CXCR2, CCR5), and LukM/LukF’ for bovine CCR2. These results highlight the importance of leukocidin-receptor interactions in pore formation and may facilitate further understanding in the role of pore-forming toxins in *S. aureus* infections. This new mechanistic insight may prove valuable to the development of future antibacterial and anti-inflammatory therapies, especially important in light of the growing menace of global antimicrobial resistance.

## MATERIALS AND METHODS

### Experimental model and subject details

#### PMN isolation, Cell Lines, and Transfections

Human blood was obtained from healthy volunteers and the polymorphonuclear (PMN) cells were isolated by Ficoll/Histopaque centrifugation [60]. Informed consent was obtained from all subjects, in accordance with the Declaration of Helsinki and the Medical Ethics Committee of the University Medical Center Utrecht (METC-protocol 07-125/C approved 1 March 2010). To ensure a truly monomeric state and prevent GFP mediated clustering of the receptor, a fusion construct of hC5aR with the monomeric GFP variant mGFP with A206K mutation (also denoted GFPmut3) [61, 62] was made at the C-terminus (primers used listed in Table 2) or a sortase A LPXTGG sequence was made in the N-terminus and cloned into pIRESpuro vectors (Table 2) by PCR. The amplification reaction was performed in three separate amplification steps using overlap extension PCR on hC5aR and mGFP templates. hC5aR (Accession number of human C5aR = NM_00173) was used as the template using enzymes and purification kits as described above. The clones were ligated into the vectors and transferred into TOP10 *E. coli* competent cells, then amplified and sequenced similarly to the toxin clones described previously. The pIRESpuro/hC5aR-mGFP vector was transfected into Human Embryonic Kidney (HEK) 293T cells (a HEK cell line, Invitrogen), stably expressing G protein Gα16, using Lipofectamine-2000 Reagent according to manufacturer’s instructions (Thermo Fisher Scientific). After 24–48 hr, transfected cells were harvested with 0.05% trypsin. To obtain a uniform, stable culture, cells were sub-cloned in a concentration of 0.5 cells/well in a 96-well plate in Dulbecco’s modified Eagle’s medium (DMEM, Lonza) supplemented with 10% fetal calf serum (Invitrogen) 100 U/ml penicillin/ 100 µg/ml streptomycin (PS; Invitrogen), 1 µg/ml Hygromycin and 250 µg/ml Puromycin. For N-terminal labelling of the sortase A recognition sequence containing HEK cells with FITC were successfully performed in two steps as described previously [63]. The expression of hC5aR was analyzed by incubating the cells in 50 µl RPMI (Invitrogen) supplemented with 0.05% human serum albumin (Sanquin), RPMI-HSA, at 5 × 10^6^ cell/ml concentration for 45 min with PE-conjugated anti-CD88 and detected by flow cytometry. The presence of mGFP or FITC-LPXTG was detected directly by flow cytometry.

**Table 2.**
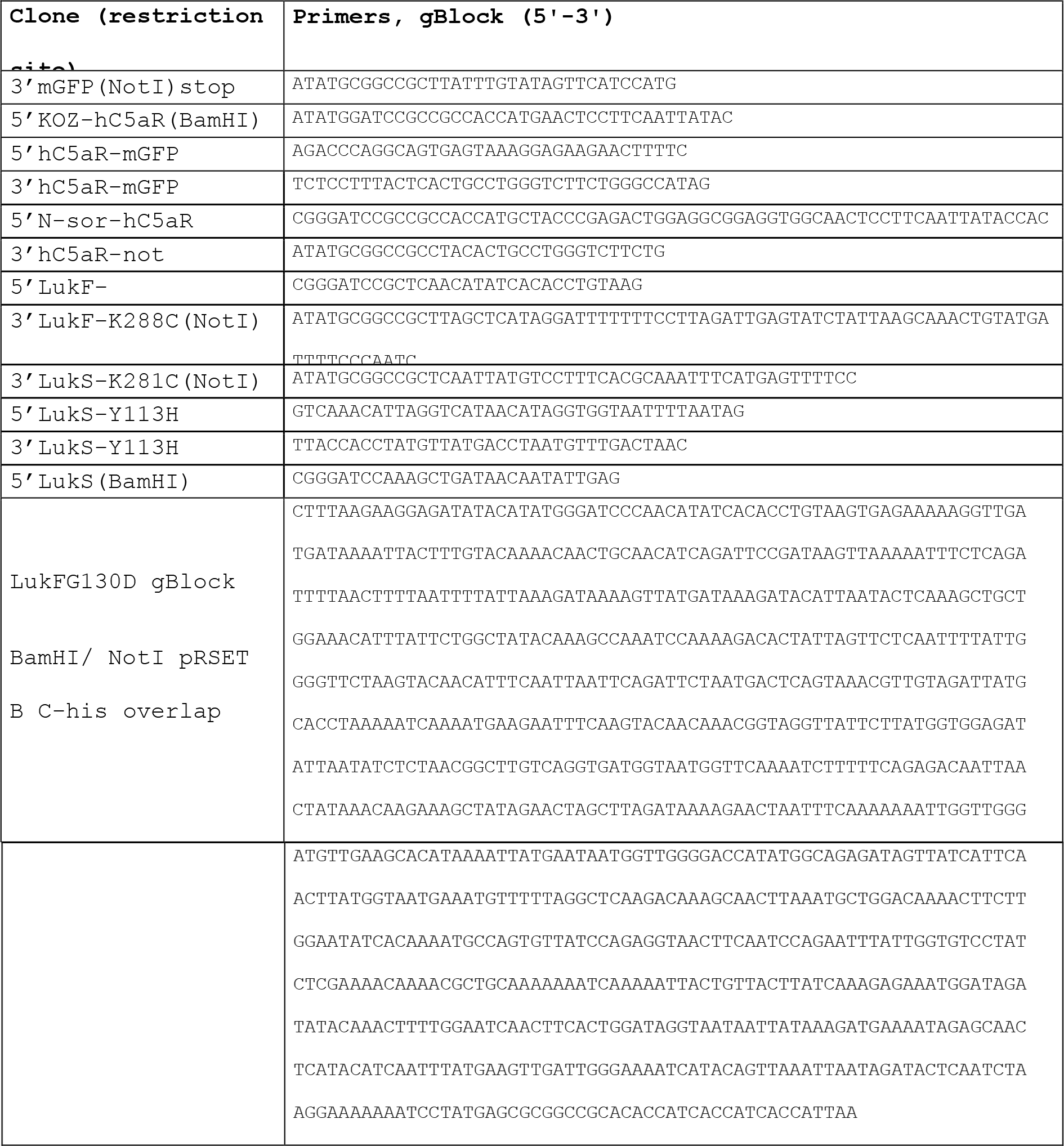
Cloning sequence details

#### Recombinant Protein Production and Purification

Polyhistidine-tagged LukS and LukF were cloned and expressed using an *E. coli* expression system. For maleimide-based labeling a single-cysteine mutation was designed to the LukS and LukF components based on previous data and the crystal structure of the octameric pore [30]. An additional mutation Y113H was included in LukS to facilitate oligomerization of the maleimide-labeled protein [17]. The target genes were amplified by PCR (Table 2) from the wild type sequences using Phusion High-Fidelity DNA polymerase (Thermo Scientific) [18]. The PCR product was cloned into a slightly modified pRSET expression vector (Invitrogen), resulting in expression of proteins with an N-terminal 6xHIS-tag. For LukF mutant G130D we useda gBlock (custom dsDNA sequence via Integrated DNA Technologies) to incorporate the LukF in the pRSET vector. Clones were sequenced to verify the correct sequence. The recombinant proteins were expressed in Rosetta Gami (DE3) pLysS *E. coli* using 1mM IPTG induction and isolated by a native isolation method. The expressed proteins were purified according to the manufacturer’s instructions (Invitrogen) using 1 ml Nickel HisTrap and Superdex 75 HiLoad columns (GE Health Care Life Sciences). Toxin components were labeled with either Cy3 (GE Healthcare), Alexa Fluor^®^ 594 or Alexa Fluor^®^ 647 C_2_ Maleimide reagent according to the manufacturer’s instructions (Thermo Scientific) resulting in negligible unlabeled content. The labeling efficiency was 100% as determined by protein concentrations using absorption at A280 and dye concentrations using absorption at A650 by a Nanodrop ND-1000 Spectrophotometer.

### Method details

#### Binding Assays

Binding of the maleimide-labeled proteins to PMN and HEK cells was confirmed by flow cytometry. LukS-K281C-Y113H (mLukS) or wild type, LukS (wt) was labeled with FITC or Alexa Fluor maleimide 647 or 594. For competition assays 3 µg/ml of the labeled protein and increasing concentration of non-labeled mLukS or LukS(wt) was incubated with isolated PMNs or HEK hC5aR-mGFP cells (5 ×10^6^ cell/ml) in a total volume of 50 µl RPMI-HSA on ice. For binding assays without competition the cells were incubated with increasing concentration of mLukF*. After 30 min incubation on ice, cells were washed, fixed with 1% paraformaldehyde and analyzed by flow cytometry. HEK cells transfected with CCR2 receptor were used as negative control for mLukS binding. To see inhibition of PE anti-CD88 (BD biosciences) binding by LukS(wt) or C5a hC5aR expressing HEK cells were first incubated with increasing concentrations of LukS or C5a for 45 min at 4°C. Then 2 µl of anti-CD88/200,000 cells was added and incubated as previously. Cells were washed once with RPMI-HSA, fixed with 1% paraformaldehyde and analyzed by flow cytometry. To detect hC5aR dissociation using sublytic concentrations of LukSF hC5aR expressing HEK cells were incubated with 100 nM of wild type LukS for 45 min at 4°C. After washing the unbound LukS sublytic concentrations of wild type LukF was added to the cells and incubated for 20 min at 37°C and 5% CO_2_ atmosphere. Percentages of lysed vs. non lysed cells were measured by using 1µg/ml of DAPI in the reaction. For C5a rebinding assay 1 µM of C5a (Sigma) was labeled with NT-647 according to manufacturer’s instructions (Monolith NT™). Free label from the sample was removed by three times centrifugation trough Amicon Ultra 0.5 mL centrifugal filters (Sigma). The hC5aR expressing HEK cells were incubated with 1 µM of wild type LukS for 45 min at 4°C. After washing 20 nM of NT647-C5a and increasing concentrations of wild type LukF was added to the cells and incubated for 20 min at 37°C and 5% CO_2_ atmosphere. Cells were washed once with RPMI-HSA, fixed with 1% paraformaldehyde and analyzed by flow cytometry. Percentages of lysed vs. non lysed cells were measured by using 1µg/ml of DAPI in the reaction. *S. aureus* Ecb (extracellular complement binding protein) was used as negative control as it interacts with another cell surface receptor, CR1 [64]. Flow cytometry data were analyzed using FlowJo v10 software package.

#### Cell Permeability Assays

Isolated PMNs or HEK hC5aR-mGFP cells (5 ×10^6^ cell/ml) were exposed to labeled and unlabeled mixtures as appropriate of mLukF/mLukS recombinant proteins at equimolar concentrations in a volume of 50 µl RPMI-HSA with 1 µg/ml of DAPI. Cells were incubated for 30 min at 37°C with 5% CO_2_ and subsequently analyzed by flow cytometry. To calculate the lysis time cells were first incubated with 150 nM of mLukS for 15 min. Then 600 nM of mLukF was added and immediately subjected to flow cytometry analysis where the permeability was measured at several time points. Cell lysis was defined as intracellular staining by DAPI. HEK cells transfected with human CCR2 receptor was used as negative control for toxin-mediated lysis. Statistical differences between means of repeated experiments were calculated using two-tailed Student *t*-tests.

#### Ex vivo complement activation assay

To maintain complement activity the blood samples were anticoagulated with lepirudin (Refludan, Schering, Berlin, Germany). Increasing concentrations of LukSF or LukS (0 to 2000 nM) was incubated in full blood for 30 min at 37°C under continuous rotation (300 rpm). Complement activity was stopped by adding 10 mM EDTA in the suspension and the plasma was separated from the blood cells by centrifugation at 5000 × g. A 1:30 dilution of each plasma sample was analyzed by SC5b 9 Enzyme Immunoassay according to manufacturer’s instructions (MicroVue SC5b 9 Plus Enzyme Immunoassay, Quidel). One *S. aureus* colony (1 x 10^8^ cells) was used as a positive control for SC5b-9 formation. The bacteria were grown over night on blood agar plate at 37°C 5% CO_2_ atmosphere.

#### Fluorescence microscopy

Cells were imaged using a Nikon A1R/STORM microscope utilizing a x100 NA oil immersion Nikon TIRF objective lens. We used a total internal reflection fluorescence (TIRF) microscopy module. We used laser excitation at wavelengths 488 nm (for mGFP), 561 nm (for Alexa594) and 647 nm (for Alexa647) from a commercial multi laser unit fiber-coupled into the microscope, capable of delivering maximum power outputs up to ~200 mW, with a depth of penetration in the range ~100-130 nm for the TIRF excitation evanescent field. Fluorescent images acquired on an iXon+ 512 EMCCD camera detector (Andor) at a magnification of 150nm/pixel. Green and red channel images were obtained by imaging through separate GFP or Alexa647 filter sets. For high laser excitation intensity single-molecule millisecond imaging, green channel images to determine mGFP localization were acquired continuously using 488 nm wavelength laser excitation over a period of ~5 min through a GFP filter set, then the filter set was manually switched to Alexa647 as for red channel images acquisition continuously using 647 nm wavelength laser excitation until complete photobleaching of the sample after 1-2 min. For photobleaching laser powers ranged between 15 mW (Alexa 647) to 100 mW (mGFP). For fixed cell analysis cells were either incubated first with mLukS or mLukS*, washed and incubated with mLukF or mLukF*, or incubated first just with mLukS* with mLukF* absent, then washed and fixed with 1% paraformaldehyde.

For fluorescence imaging the HEK cells were grown on 0.1% poly-L-lysine which coated 8-well chambered cover glass slides (Ibidi) in standard growth conditions described above. To analyze the deposition of mLukS* on live cells the cells were first imaged in PBS buffer in the absence of toxin. Here, a 256 × 256 pixel area covered approximately one cell per field of view. Then the cells were incubated for 2 min with 5 µg/ml of Alexa594 maleimide-labeled LukS in RPMI-HSA and the cells were carefully washed with PBS keeping the imaging area and focus constant. Because of the fast bleaching of the Alexa647 label a more stable Alexa594 label was used for the LukS deposition imaging. The deposition of mLukS was detected for 10 min and the lysis of the cell was recorded for 15 min after addition of 600 nM of unlabeled mLukF. Cells were imaged in TIRF at 50 ms per frame with the laser automatically switched between 488 nm/0.22 mW, 647 nm/3 mW and 561 nm/3 mW or 488 nm/0.22 mW and 561 nm/3 mW (Figure 2).

#### RFRET experiments

The Cy3-labeled LukS or non-labeled LukS and Cy3-labeled LukF or non-labeled LukF or Cy-labeled LukF and controls (only FITC-labeled cells or unlabeled cells with only Cy3-labeled LukS) were incubated at 4°C for 30 min and then 10 min at 37°C, washed 2 x with RPMI-HSA and fixed as before. The cells were in PBS during imaging. FITC experiments were performed using Leica TCS SP5 microscope, using a 62× oil immersion objective lens, and FRET Sensitized Emission Wizard in Leica Application Suite Advanced Fluorescence (LAS AF). Images were acquired using 488 nm and 543 nm wavelength lasers and a laser power of 27% 12.0 (A.U.) and a scan size of 512 × 512, 800 ms, 50 ms per frame, beam splitter TD 488/543/633.

FRET efficiency ε was calculated using the donor, directly excited acceptor and donor excited acceptor intensity from n=5-10 manual regions of interest inside cells for each experiment, using the following formula [65]:

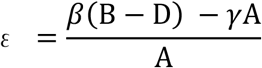

B = Intensity signal, donor excited acceptor

D = Intensity signal, donor

A = Intensity Signal, directly excited acceptor

β = calibration for ratio of measured intensities of B_donor channel_/A_donor channel_

γ = calibration for ratio of measured intensities of B_acceptor channel_/A_acceptor channel_

#### Single-molecule imaging of live and fixed cells

GFP and Alexa647 fluorescence micrograph time series of fixed and live cells were sampled taken at. 20 ms per frame. Green channel images were acquired continuously using 488 nm wavelength laser excitation over a period of *ca*. 5 min via the GFP filter set. Then the filter set was manually switched to that for Alexa647 and red channel images were acquired continuously using 647 nm wavelength laser excitation was until complete photobleaching of the sample after 1-2 min. The step-wise single-molecule fluorescence photobleaching was analyzed both for live and fixed cells. For live cell photobleaching analysis the cells were incubated with 150 nM of labeled or unlabeled mLukS as required for *ca*. 15 min. After washing with PBS, 600 nM of labeled or unlabeled mLukF was added and the imaging was done immediately within 10-15 min. If labeled LukF was added the wells were washed with PBS before analysis. Also samples with only LukS and without toxins were analyzed. For fixed cell analysis the cells were incubated first with mLukS or mLukS* for 30 min at +4°C in RPMI-HSA, washed with same buffer and incubated for 10 min at 37°C with mLukF or mLukF*, or the same protocol was followed but using mLukS* alone with mLukF* absent. Then the cells were washed and fixed with 1% paraformaldehyde. 1M mercaptoethylamine (MEA) buffer was used for fixed cell analysis. Photobleaching of recombinant mGFP and mLukS-Alexa647 were also separately analyzed in a tunnel slide comprising two pieces of double-sided tape forming a channel sandwiched between a standard glass microscope slide and a plasma cleaned coverslip. Proteins solutions (1μg/ml) were immobilized onto the coverslip coated by anti-GFP or anti-His antibodies respectively with phosphate buffered saline washes in between.

### Quantification and statistical analysis

#### Binding and permeability assays

Statistical significance between repeated (n>1) experiments was analyzed using two-tailed Student’s *t*-tests where using a standard p-value threshold of <0.05 as indicating statistical significance. Means and standard deviations of repeated experiments are shown in error bars, unless indicated otherwise.

#### Image analysis

Basic image extraction, cropping and quantification was done using NIS-Elements microscope imaging software and Image J. More advanced foci tracking was done using bespoke software written in MATLAB (Mathworks) [36] which enabled automatic detection and localization of individual fluorescent foci to within 40 nm lateral precision (Additional file 2: Figure S2a). The software identifies candidate foci by a combination of pixel intensity thresholding and image transformation. The intensity centroid and characteristic intensity, defined as the sum of the pixel intensities inside a 5-pixel radius circular region of interest around the foci intensity centroid minus the local background and corrected for non-uniformity in the excitation field are determined by repeated Gaussian masking. If the signal-to-noise ratio of a foci (the intensity per pixel/background standard deviation per pixel) is greater than a pre-set threshold, nominally here set at 0.4 based on prior simulations, it is accepted and fitted with a 2D radial Gaussian function to determine its width. Foci in consecutive frames within a single PSF width, and not different in intensity or width by greater than a factor of two, are linked into the same track.

Foci intensity was used to quantify stoichiometry information. As foci photobleach over time during continuous laser excitation their intensity falls in a stepwise manner due to photobleaching of an integer number of fluorophore tags in each sampling time window. By quantifying the size of a single step, the characteristic intensity of a single fluorophore can be obtained and thus the stoichiometry of the foci from its initial intensity. The step size is found from the periodicity in the distribution of foci intensities corroborated by the pairwise distance (PwD) distribution of these intensities and the Fourier spectrum of the PwD which contains peaks at the characteristic intensity and harmonics at multiples of this value (Additional file: Figure S2d-e).

Here, the copy number of hC5aR-mGFP was comparatively high such that the TIRF images were initially saturated in regards to pixel intensity output. After ~20 s of photobleaching the non-saturated foci intensity values were fitted by an exponential function which characterized the rate of intensity decay, equivalent to an exponential photobleach time of ~20 s, and extrapolated back to zero time to determine the initial foci intensity (Additional file: Figure S2f). The Alexa647 dye also bleached during 647 nm wavelength laser excitation but images were not initially saturated, but also in some images which were exposed to the 488nm laser and then the 647nm laser, also bleached by the 488 nm wavelength laser. In these images a fixed correction factor of 6x, determined by comparing to images exposed to the 647 nm laser first, was used. The stoichiometry of each foci was then determined as the initial intensity divided by the intensity of the appropriate single fluorescent dye tag (i.e. either mGFP or Alexa647 in this case).

We characterized the mobility of tracked foci by calculating their MSD as a function of time interval (τ). For each detected foci the MSD was calculated from the measured intensity centroid (*x*(*t*),y(*t*)) at time *t* assuming a foci track of *N* consecutive image frames at a time interval τ=*n*Δ*t* where *n* is a positive integer and Δ*t* is the frame integration time (here 20 ms):

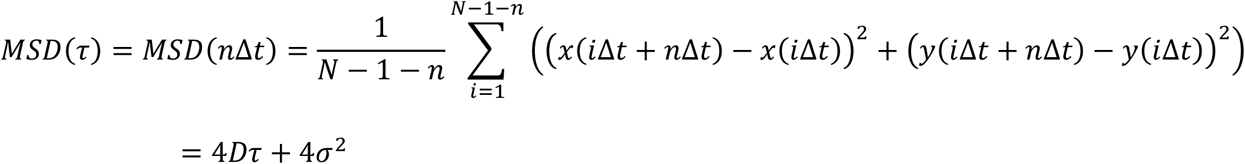

The lateral (*xy*) localization precision is given by *σ* which we determine to be 40 nm. We fitted a straight line to each separate MSD relation. Assuming a line fit has an optimized gradient *g* to the first 4 points (defined as the first 3 measured MSD data points for *n*=1, 2 and 3, in addition to a 4^th^ data corresponding to *n*=0 obtained from constraining the intercept be 4*σ*^2^ to within measurement error of the localization precision) then estimated the microscopic diffusion coefficient *D* as *g*/4.Δ*t*. For immobile foci, tracks were collated and compiled to generate a mean MSD vs. τ relation which was fitted to an asymptotic rising exponential function as an analytical model for confined diffusion of MSD plateau equal to *L*^2^/6 where *L* is the effective confinement diameter [42], enabling us to estimate the confinement diameter.

#### Colocalization analysis

The extent of colocalization between red and green detected foci was determined using a method which calculated the overlap integral between each green and red foci pair, whose centroids were within ~1 PSF width (~3 pixels). Assuming two normalized, two-dimensional Gaussian intensity distributions *g_1_* and *g_2_*, for green and red foci respectively, centred around (*x_1_*, *y_1_*) with sigma width *σ_1_*, and around (*x_2_*, *y_2_*) with width *σ_2_*, the overlap integral ν is analytically determined as:

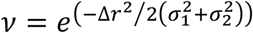

where:

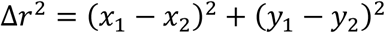

We use a criterion of an overlap integral of 0.75 or above to indicate putative colocalization [41] since this corresponds to a foci centroid separation equivalent to the localization precision in this case. By quantifying the standard deviation on the number of detected foci in each channel we estimate that the standard error of colocalisation proportion under our typical imaging conditions is approximately 9%.

#### Random foci overlap models

We calculated the probability of foci overlap in a single color channel by first estimating a sensible range of foci surface density *n*. For the lower limit we used the number of foci tracks detected in a 20 image frame time window, for the upper limit we used the average measured value of the background-corrected pixel intensity value divided by the intensity of a single fluorophore (equivalent to ~1 mLukS* molecule per pixel). We implemented these probability estimates into a surface density model which assumed a random Poisson distribution for nearest-neighbor separation [40, 41, 66-69]. This model indicates that the probability that a nearest-neighbor separation is greater than *w* is given by exp(-π*w*^2^*n*). The probability of overlap for each density estimate (Additional file 4: Figure S4) was convolved with a real molecular stoichiometry distribution and a Gaussian function *p(x)* of stoichiometry (*x*):

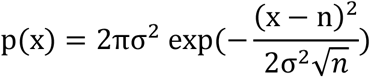

where σ is the width of single fluorophore intensity distribution (~0.7 molecules), and *n* is the real molecular stoichiometry. The tetramer model assumes *n*=4, then all higher order stoichiometries are due to overlapping PSFs. The tetramer oligomer molecule assumed an equal number of multimerized tetramers up to 5, which gave the best fit to the data.

The same strategy was used to model the random overlap probability for green and red color channel fluorescent foci in dual color imaging experiments to assess the extent of apparent colocalization due to random overlap between hC5aR and mLukS*/F*. The probability that a nearest-neighbor separation is greater than *w* for foci of two different types is the same as a single type multiplied by 2/3. (38)

### Software

All our bespoke software developed is freely and openly accessible via https://sourceforge.net/projects/york-biophysics/ (65).

### Statistical tests and replicates

All statistical tests used are two-tailed unless stated otherwise. For single-molecule TIRF imaging each cell can be defined as a biological replicate sampled from the cell population. We chose sample sizes of 5-7 cells yielding thousands of foci, generating reasonable estimates for stoichiometry and diffusion coefficient distributions. Technical replicates are not possible with the irreversible photobleaching assay.

### Ethics statement

Human polymorphonuclear (PMN) cells, obtained from healthy volunteers were isolated by Ficoll/Histopaque centrifugation [60]. Informed written consent was obtained from all subjects, in accordance with the Declaration of Helsinki and the Medical Ethics Committee of the University Medical Center Utrecht (METC-protocol 07-125/C approved 1 March 2010).

## Acknowledgments

We thank Piet Aerts and Angelino Tromp for assistance in sample preparation and labeling and Esther van’t Veld and Richard Wubbolts (Utrecht) for assistance with light microscopy and Dr Karin Strijbis (Veterinary School, UU) providing the sortase A enzyme. This work was supported by The Finnish Cultural Foundation (grants 00131060 and 00142390), the Biological Physical Sciences Institute, Royal Society, MRC (grant MR/K01580X/1), BBSRC (grant BB/N006453/1), the EPSRC Physics of Life UK network and the Wellcome Trust [ref: 204829] through the Centre for Future Health (CFH) at the University of York, UK.

## Conflict of interest

All the authors declare that they have no conflict of interests.

---

**Additional file 6: Movie S1 (.avi).** Deposition of mLukS on hC5aR-mGFP HEK cells. The movie is shown in two clips before (no toxin) and after addition of Alexa 595 labeled mLukS (add mLukS*). This movie was recorded for *ca*. 13 min and displayed here at 100x speed.

**Additional file 7: Movie S2 (.avi).** Lysis of hC5aR-mGFP HEK cells incubated with Alexa594 labeled mLukS* and mLukF. The cells were preincubated with mLukS* and the lysis of the cells were monitored for *ca*. 13 min after addition of mLukF (add mLukF). The red arrow points to the vesicles released during cell lysis. This movie was recorded for *ca*. 13 min and displayed here at 100x speed.

**Additional file 8: Movie S3 (.avi).** Imaging live hC5aR-mGFP cells. After 1-2 min of exposure, several distinct, mobile, circular fluorescent foci in the planer membrane regions were observed. Movie is displayed in real time.

**Additional file 9: Movie S4 (.avi).** Imaging mLukS* incubated with hC5aR-mGFP cells. Several distinct, mobile, circular fluorescent foci were observed. Movie is displayed in real time.

**Additional file 1: Figure S1.**
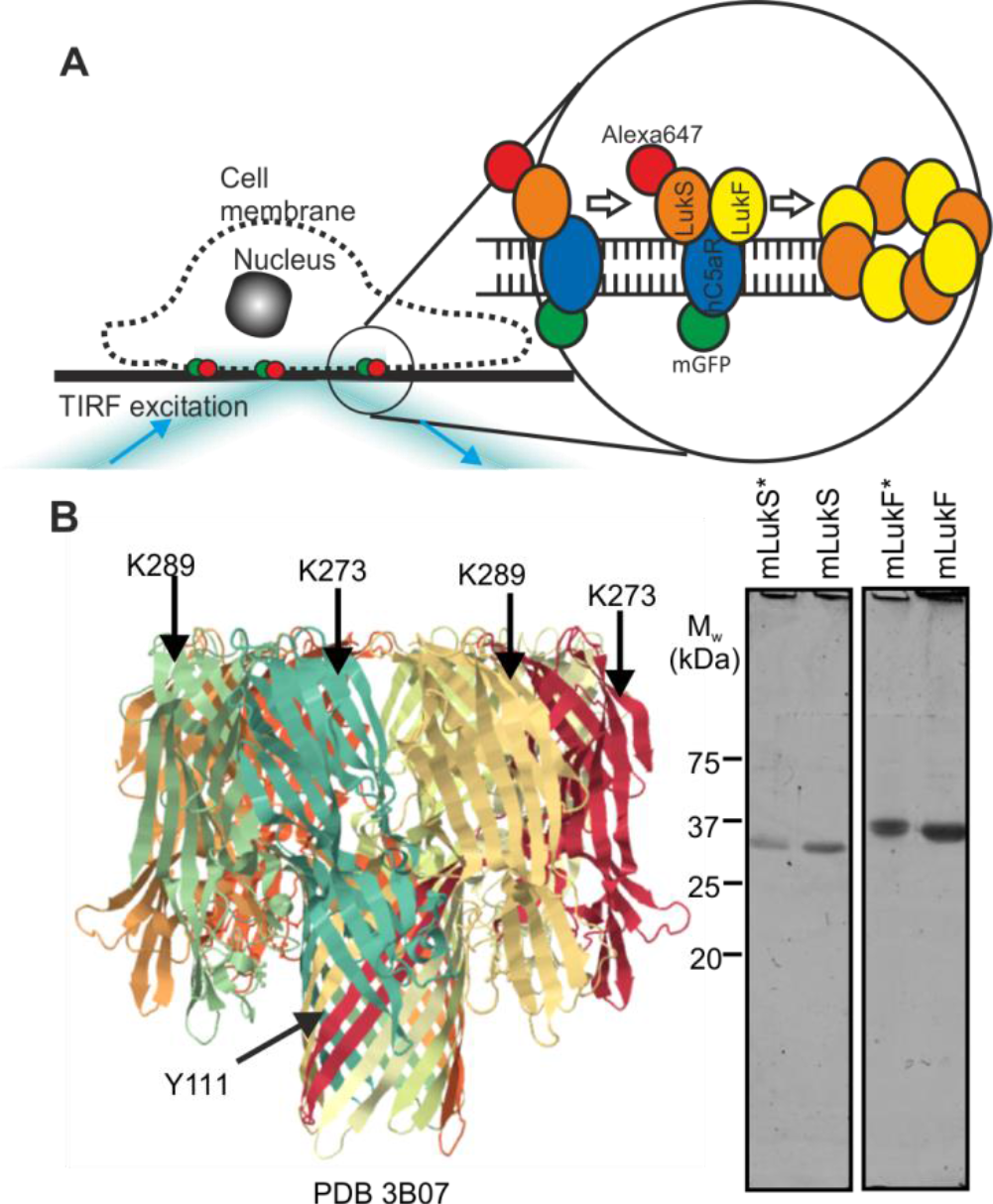
Construction of recombinant leukocidin proteins. (A) Schematic of TIRF imaging assay. (B) (left panel) Crystal structure of octameric pore complex of γ-hemolysin (PDB ID:3B07). K273/Y111 and K289 on S and F components of γ-hemolysin corresponds to our engineered mutations, K281C/Y113H and K288C, on LukSF marked in their equivalent places on γ-hemolysin; (right panel) SDS-PAGE of the unlabeled LukSK281CY113H and LukFK288C (mLukS and mLukF) and Alexa-labeled mLukS* and mLukF* toxin components, bands visible at locations consistent with molecular weight of 33 kDa and 34 kDa for LukS and LukF respectively.

**Additional file 2: Figure S2.**
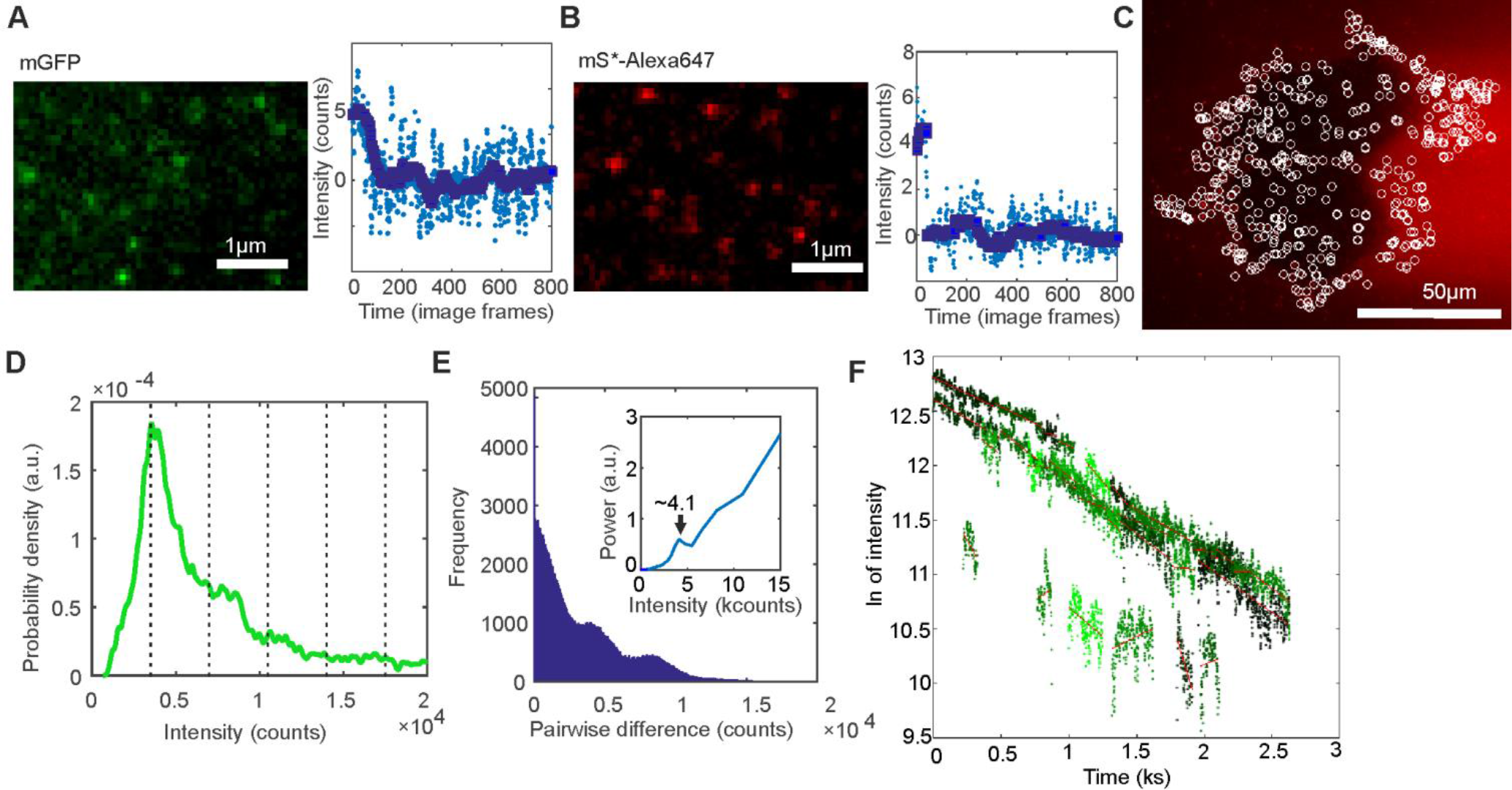
Fluorescent protein characterization. A. Fluorescence micrograph of immobilized mGFP and an intensity *vs*. time trace for one foci showing a single photobleach step. Raw data in light blue and edge-preserving Chung-Kennedy filtered data in dark blue. B. As A for LukS-Alexa647, C. LukS-Alexa647 micrograph (red) with found foci indicated as white circles. D. Intensity distribution of Alexa 647 foci intensities from whole photobleach experiment showing periodicity at ~3,500 counts on our camera detector. E. Pairwise distance distribution of intensity in D with Fourier spectrum (inset) showing peak at ~4,000 counts. F. GFP foci intensity (natural log) time traces (green) with linear fits (red).

**Additional file 3: Figure S3.**
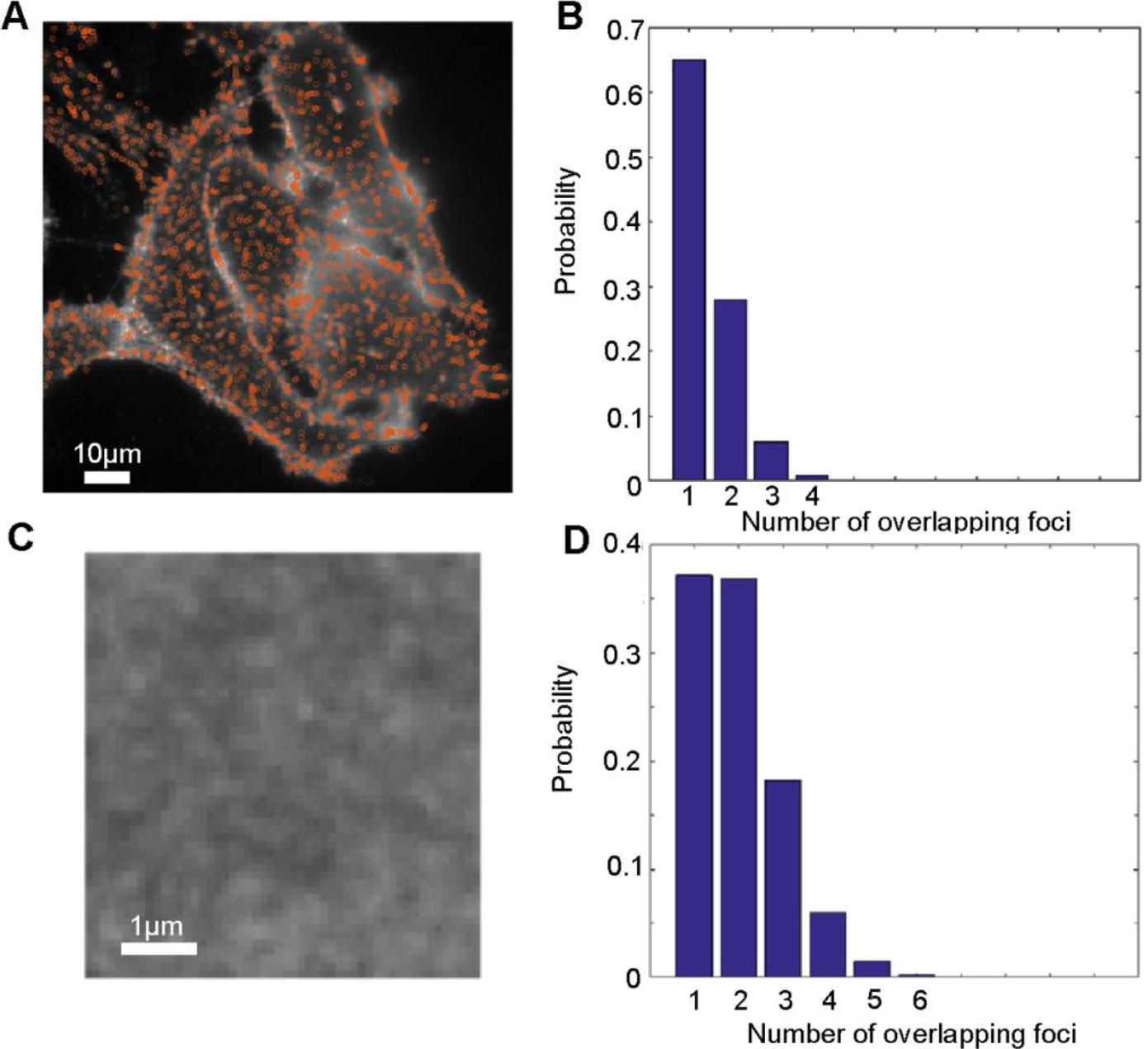
Density of LukS spots. A. micrograph of mLukS* (white) with found foci (orange circles) B. Probability distribution of overlap frequency using spots in A to calculate density. C. Zoom in of mLukS micrograph. D. Probability distribution of overlap frequency using intensity in C to calculate maximum density estimate.

**Additional file 4: Figure S4.**
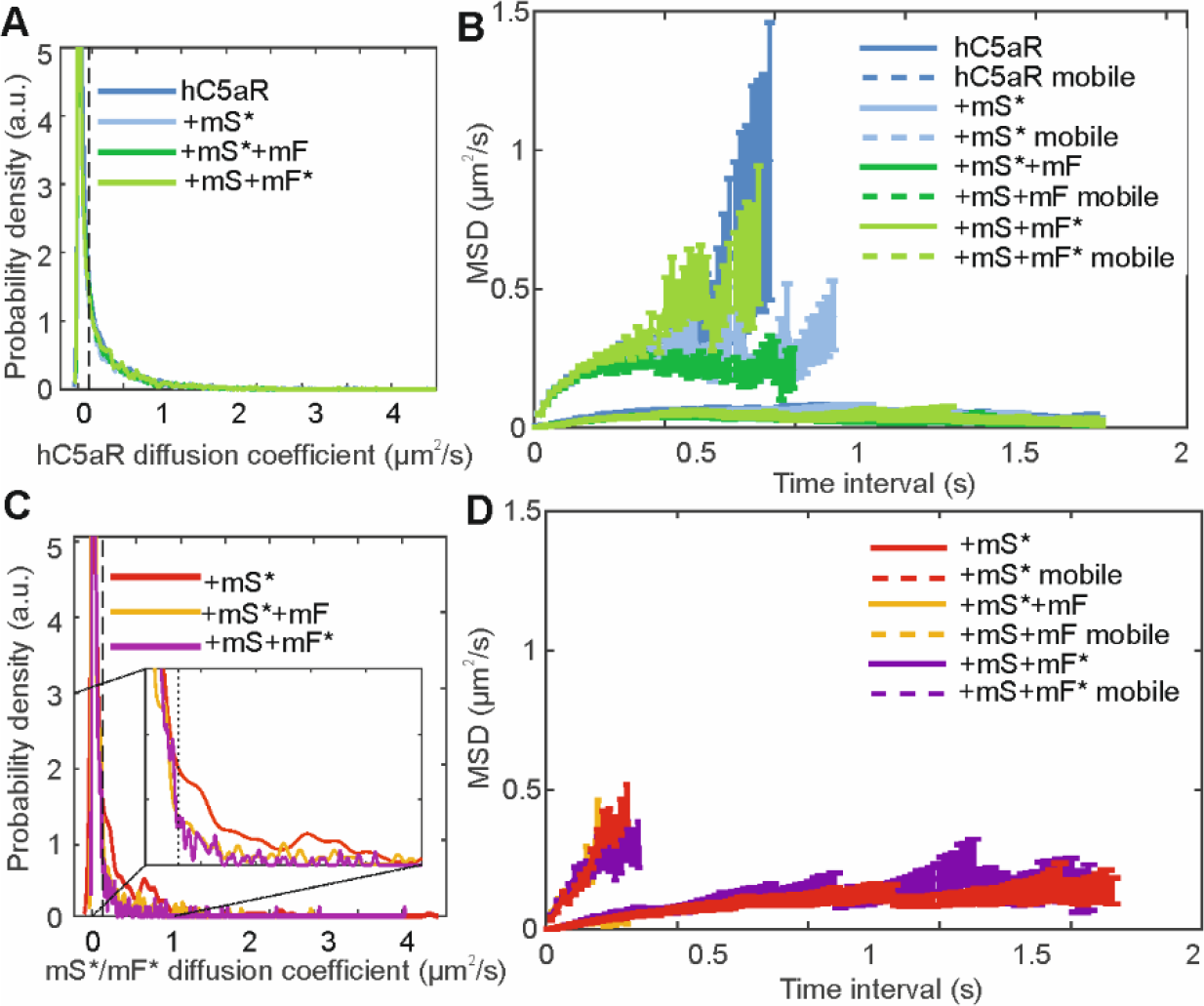
Mobility analysis. A. The probability distribution of microscopic diffusion coefficient showing the threshold for immobility as black dotted line and B. the mean squared displacement against time interval for mobile (upper) and immobile (lower) of hC5aR. C. and D. similar for mLukS/F*. An insert showing a zoomed in portion of the plot is shown in C. to better illustrate the division between mobile and immobile.

**Additional file 5: Figure S5.**
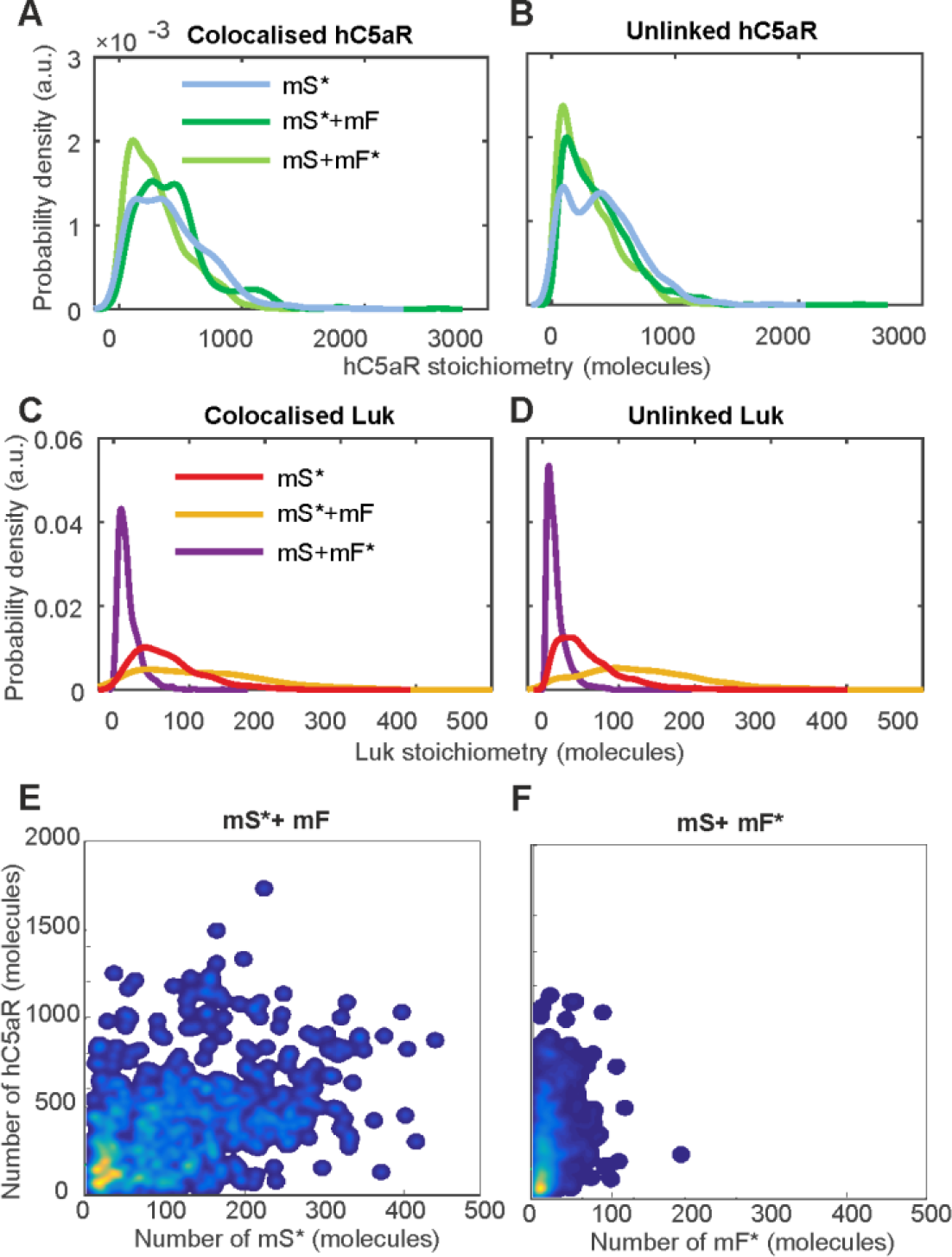
Colocalization analysis. The probability distribution of linked (A) and unlinked (B) hC5aR and similar for mLukS* (C. and D.). E. and F. False-color heatmap scatter plots indicating that h5CaR stoichiometry is uncorrelated to mLukS or mLukF stoichiometry in the presence of mLukF.

